# BRN2 and PTN unveil multiple neurodevelopmental mechanisms in Schizophrenia patient-derived cerebral organoids

**DOI:** 10.1101/2021.06.10.447949

**Authors:** Michael Notaras, Aiman Lodhi, Friederike Dundar, Paul Collier, Nicole Sayles, Hagen Tilgner, David Greening, Dilek Colak

## Abstract

Due to an inability to ethically access developing human brain tissue as well as identify prospective cases, early-arising neurodevelopmental and cell-specific signatures of Schizophrenia (Scz) have remained unknown and thus undefined. To overcome these challenges, we utilized Scz patient-derived stem cells to generate 3D cerebral organoids to model neuropathology of Scz during this critical period. We discovered that Scz organoids exhibited ventricular neuropathology resulting in altered progenitor survival and disrupted neurogenesis. This ultimately yielded fewer neurons within developing cortical fields of Scz organoids. Single-cell sequencing revealed that Scz progenitors were specifically depleted of neuronal programming factors leading to a remodeling of cell-lineages, altered differentiation trajectories, and distorted cortical cell-type diversity. While Scz organoids were 99.95% similar in their macromolecular diversity to Ctrls, four GWAS factors (PTN, COMT, PLCL1, and PODXL) and peptide fragments belonging to the POU-domain transcription factor family (e.g. POU3F2/BRN2) were altered. This revealed that Scz organoids principally differed not in their proteomic diversity, but specifically in their total quantity of disease and neurodevelopmental factors at the molecular level. Single-cell sequencing also subsequently identified cell-type specific alterations in neuronal programming factors and growth factors, and specifically replicated the depletion of POU3F2 (BRN2) and PTN in both Scz progenitors and neurons. Consequently, in two mechanistic rescue experiments we identified that the transcription factor POU3F2 (BRN2) and growth factor PTN operate as mechanistic substrates of neurogenesis and cellular survival, respectively, in Scz organoids. This suggests that multiple mechanisms of Scz exist in patient-derived organoids, and that these disparate mechanisms converge upon primordial brain developmental pathways such as neuronal differentiation, survival, and growth factor support, which may amalgamate to elevate intrinsic risk of Scz.

## INTRODUCTION

Schizophrenia (Scz) typically emerges in early adulthood and is a chronic brain disorder that affects ∼1% of the population. Upon onset, Scz neuropathology is already sufficiently vast that mechanistic studies have failed to pinpoint definitive causes of disease, as well as whether hallmark pathology are *causes* or *consequences* of disease progression. While adolescent brain development is established as a critical period for Scz onset [1], Scz appears to maintain earlier developmental origins [2]. Due to ethical and technical constraints that prevent the collection of biological specimens at early developmental time-points, pre-disease signatures of Scz prior to adolescence remain unknown. Thus, defining the timing of developmental neuropathology in Scz, as well as the temporally specific involvement of putative biological risk factors, could lead to the identification of common pathways regulating ontogenesis of disease.

Evidence that Scz risk may commence *in utero* has been collected for decades now, and epidemiological evidence suggests that prenatal stress [3], prenatal famine [4], early vitamin D deficiency [5], as well as maternal immune challenges [6, 7] may elevate Scz risk. Indeed, risk may manifest as early as the first trimester of brain development [3].

However, little is known of the mechanisms involved in prenatal Scz risk, and almost all knowledge of Scz developmental phenotypes have emerged from maternal immune activation (MIA) models of Scz. Modeling MIA has subsequently unveiled alterations in tonic neurotrophic support *in vivo* (e.g. BDNF and TrkB expression) [22], perturbations in neuronal differentiation of progenitors *in vitro* [23], as well as effects on new-born neuron birth *in vivo* [24]. Additionally, MIA Scz models have reported increased cell death [20] and delayed axon development [21]. These phenotypes are of particular noteworthiness given that similar alterations have been observed in Scz postmortem studies. This includes alterations in growth factors (e.g. BDNF, NT-3, and NT-4) [25–27], neurotrophin receptors [28, 29], cell death markers [30], neural stem cell proliferation [31], and reduced new-born cell numbers [32]. Together, these animal and human studies implicate that factors and pathways related to brain development may regulate Scz risk.

Genetic studies have also converged upon brain development factors as loci of Scz risk. Analysis of tens of thousands of Scz patients has revealed at least 108 prominent loci of risk [36], including previously unknown regions that may encode novel development and growth factors (e.g. Pleiotrophin, otherwise known as PTN [36]). However, many association studies have also produced inconsistent linkage (for e.g., the *BDNF* gene locus, see discussion in [33–35]). Therefore, mechanisms of Scz have remained not just elusive, but also speculative.

To overcome these gaps, here we generate patient-derived cerebral organoids to investigate early developmental pathology of Scz in 3D human-derived tissue. We found that Scz cerebral organoids exhibited abnormalities in ventricular zone progenitors, which died at increased rates and also exhibited disrupted neuronal differentiation.

There was a corresponding depletion of early-born, late-born, and pan (MAP2+) neuron numbers in 3D cortical assemblies of Scz organoids. In a search for the molecular underpinnings of these neurodevelopmental features, we adapted unbiased mass-spectrometry and found that Scz organoids differed not in their molecular diversity, but rather in their expression of development factors (POU-domain transcription factors, e.g. BRN2) as well as specific Scz risk factors (e.g. PTN). Single-cell RNA-sequencing (scRNA-Seq) confirmed that Scz organoids were depleted of progenitors and neurons, principally due to remodeling of progenitor cell-lineages yielding altered differentiation trajectories. Additionally, scRNA-Seq confirmed the depletion of *BRN2* and *PTN* gene-expression in Scz progenitors and neurons. In separate experiments, we modulated BRN2 or PTN levels in Scz organoids and both factors rescued neuron numbers, albeit via seemingly different mechanisms. In sum, high-content analysis revealed cell-specific neuropathology and multiple mechanisms of Scz within 3D patient-derived organoids, which unveiled distinct developmental pathways and novel disease factor mechanisms that define Scz risk during early corticogenesis.

## RESULTS

### Patient-derived Scz organoids exhibit ventricular entropy and neuropathology

To study neurodevelopmental aspects of Scz we generated self-assembling, self- maturating, and human-derived cerebral organoids [37]. Cerebral organoids recapitulate transcriptomic [38] and epigenomic [39] programs of fetal development that drive the assembly of ventricular progenitor zones that give rise to neuronal ensembles [7, 8]. This broadly mimics the first trimester of human brain development [37, 40]. Recent organoid models often apply exogenous growth factors (e.g. BDNF [44–46], NT-3 [46], and EGF [46]) and/or pathway modulators (e.g. TGFβ and WNT pathway inhibitors [47, 48]) to forcibly direct the differentiation of progenitors towards a neuronal fate or promote their survival by reducing apoptosis. However, factors such as BDNF and related neurotrophins [25–27, 34], TGFβ [49–54], and WNT signaling [55–60] as well as other morphogens are longstanding contributors to Scz pathophysiology and may play a role in the ontogenesis of disease during early neurodevelopment. Indeed, these same factors and pathways have hypothesized roles in early disease development via their effects on stem cells and neuronal programming [61]. Therefore, given our particular interest in disease modeling, we adapted a morphogen-free organoid protocol [37] for our experiments to avoid these potential confounding factors.

We consequently generated 3D cerebral organoids from healthy controls (Ctrls; who had no personal history nor first-degree relative with a psychiatric diagnosis) and idiopathic Scz cases (for clinical notes, see Supplementary Table 1). Cerebral organoids generated from both Ctrl and Scz patients were subjected to rolling quality control assessments (see Methods) and all iPSC lines exhibited robust evidence of neural induction and little variability in progenitor numbers, death, mitotic activity, and total neuron numbers at baseline (see fig. S1). This included the formation and presence of ventricles and ventricular zones that were comprised of neural progenitors (SOX2+, and NESTIN+ progenitor cells) that produced extensive neuronal fields (filled with β3+ and MAP2+ neuronal cells). Similar to Ctrl organoids, Scz organoids also exhibited extensive neural induction that was defined by the presence of NESTIN+ fibers and SOX2+ progenitor-enriched ventricular zones (see fig. 1b). However, relative to Ctrls, Scz organoids exhibited ventricular zones that appeared smaller than Ctrls and cortical fields that were more sparsely filled with neurons. These phenotypes mutually suggest that Scz progenitors may be prematurely dying, or unable to initiate neurogenesis. An analysis of progenitor cell death in a large number of iPSC donors (*n* = 21; Ctrl = 4-5 donors and Scz = 15-17 patients) revealed that Scz progenitors exhibited both a significant increase in ventricular progenitor cell death (fig. 1c) and a significant decrease in the number of neurons within Scz cortical fields. In an independent, pseudorandomly sampled cross-sectional cohort, increased apoptosis was replicated in Scz organoids via three independent markers of cell death induction (increased DNA fragmentation, pH2AX+ DNA damage, and PARP+ cells, see fig. S2). This suggests that due to increased ventricular progenitor death, fewer Scz progenitors are able to successfully differentiate into neurons.

**Figure 1.**
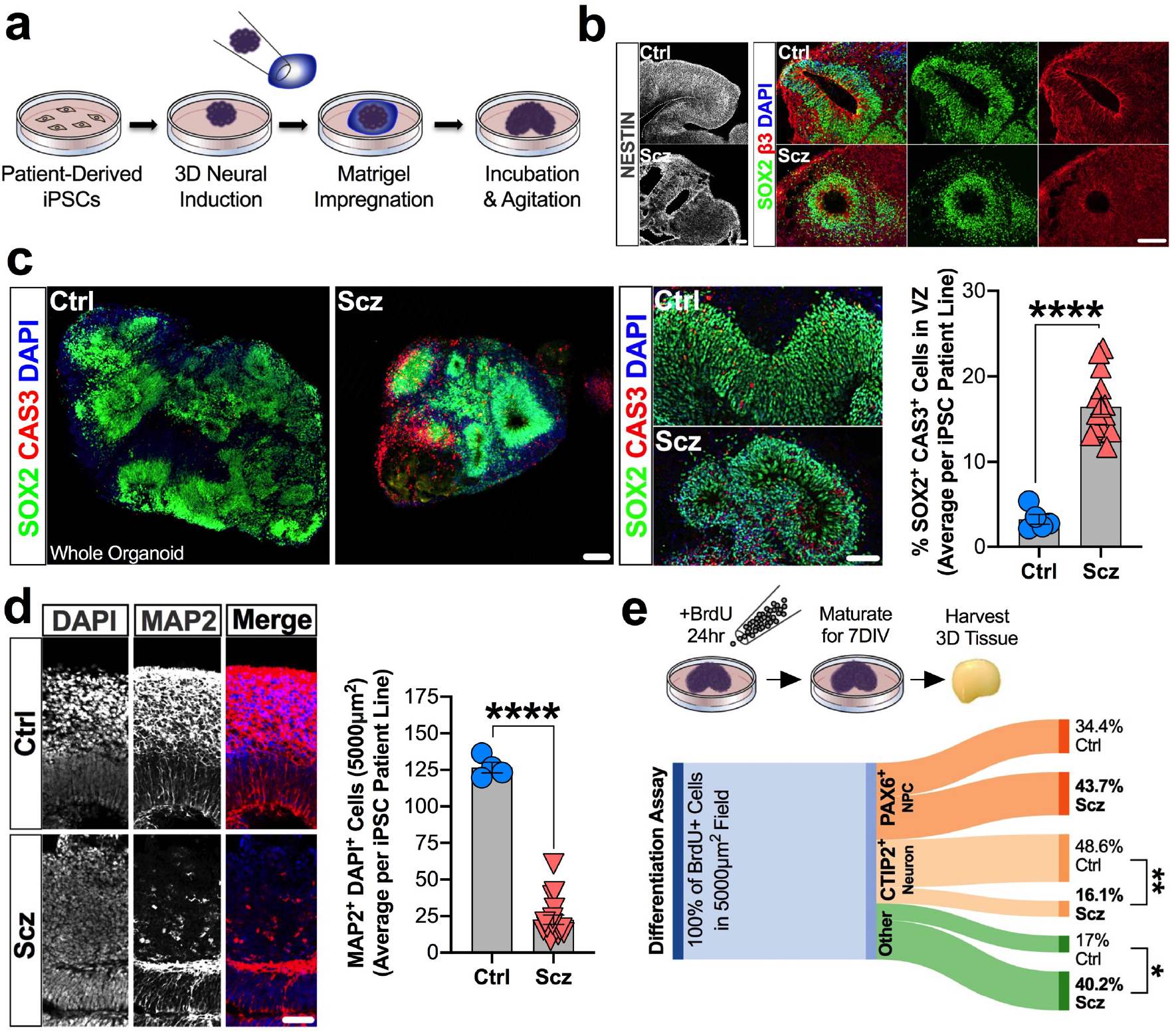
Ventricular progenitors are disrupted in Scz patient-derived organoids. a, Schematic of 3D cerebral organoid generation pipeline. Briefly, two-dimensional iPSC colonies were dissociated, cultured into 3D embryoid body aggregates, differentiated towards a neuroectodermal fate, embedded in a Matrigel droplet to provide a matrix for expansion, and maturated upon an orbital shaker. **b, Representative images of neural induction in 3D cerebral organoids.** Shown is a representative image of a single ventricular zone from each group, exhibiting filamentous processes of neural stem cells (NESTIN+ cells), ventricular neural progenitors (SOX2+ cells), and their radially organized fibers which lead to dense neuronal fields (β-tubulin III; β3).**c, Scz ventricular zone neural progenitors exhibit increased apoptosis.** Analysis of co-localization of cleaved (activated) CAS3+ with SOX2+ cells revealed that neural progenitor apoptosis was significantly elevated in Scz organoids compared to Ctrl organoids (total *n* = 20 iPSC donors; Ctrl *n* = 5 and Scz *n* = 15). Scz organoids exhibited increased progenitor death irrespective of ventricular zone size and exhibited stable rates of error across ventricular zones (data not shown). In follow-up orthogonal validation experiments, increased death was replicated in an independent cohort via single-cell analysis of DNA fragmentation, DNA damage induction, and expression of cleaved PARP (see fig. S2). Together, these data establish progenitor entropy as an intrinsic Scz phenotype in cultures of 3D patient-derived neural tissue. **d, Schematic of progenitor neuropathology in Scz organoids.** Cortical development is defined by the sequential differentiation of neural progenitors into neurons that reside within the developing cortical plate. Due to a disruption in the survival of ventricular progenitors, we next assessed neuron numbers in Ctrl and Scz organoids. Scz organoids exhibited a decrease in the numbers of MAP2+ neurons relative to Ctrls (total *n* = 21 iPSC donors; Ctrl *n* = 4 and Scz *n* = 17). e, Neuronal differentiation is disrupted in Scz organoids. Lastly, we sought to determine if disrupted differentiation also contributes to neuronal depletion in Scz organoids. To label dividing progenitor cells and assess their fate, we used a 24hr BrdU pulse and 7d chase paradigm in a pseudorandomly selected cross-sectional cohort. Data is provided as a Sankey flow visualization so that differentiation can be evaluated as the proportion of all BrdU+ cells within a field. Approximately half of the BrdU+ cells differentiated into CTIP2+ early-born cortical neurons (48.6%) in Ctrl organoids. Contrary to this, Scz organoids exhibited a disrupted developmental trajectory whereby only 16.1% of cells differentiated into early-born CTIP2+ neurons (total *n* = 8 iPSC donors; *n* = 4 Ctrl and *n* = 4 Scz). While a trend for increased self-renewal of PAX6+ progenitors was detected in Scz organoids (43.7% BrdU+ cells) relative to Ctrls (34.4% BrdU+ cells), this comparison was not significant. Thus, in addition to progenitor death, Scz organoids exhibit fewer neurons in developing cortical fields due to a mechanism that restrained neurogenesis by altering the differentiation trajectory of neural progenitors. \**p* < 0.05, \*\*\*\**p* < 0.0001. Error bars reflect Standard Error of the Mean. Scale bar: b and d = 60µm, whole organoid panel in c = 60µm, far right panel of c = 100µm. Ctrl: Control, Scz: Schizophrenia, VZ: Ventricular Zone.

However, these data do not rule out the possibility that Scz progenitors that were the not subject of premature death may also exhibit a difference in their neurogenic potential.

That is, while progenitor cell death as likely to be an important rate-limiting factor in the upstream depletion of neurons within Scz organoids, it was also possible that other mechanisms may simultaneously restrain new-born neuron generation in Scz organoids. To resolve this, we adapted a BrdU pulse-chase paradigm that comprised 24hr BrdU pulse and a 7d chase period (fig. 1e) that would identify if an intrinsic difference in the proportion of new-born neurons being generated also co-existed within Scz organoids.

Following our pulse-chase regime, we harvested organoids and examined the proportion of BrdU+ proliferating progenitors that underwent neuronal differentiation (BrdU+ CTIP2+) or progenitor self-renewal (BrdU+ PAX6+) within the same field. This allowed us to map neuronal cell fate decisions as well as determine whether Scz progenitors were preferentially self-renewing instead of generating neurons. In Ctrl organoids, progenitors predominantly differentiated into early-born neurons within our 7d chase period. However, Scz organoids exhibited significantly disrupted neurogenesis relative to Ctrls (48.6% vs 16.1% in Scz organoids). In sum, these data revealed that neuronal loss in Scz organoids was found to arise from an amalgamation of *both* elevated apoptosis and decreased neurogenesis.

### Proteome mapping reveals enrichment for disease factors in Scz organoids

Having identified two cellular phenotypes (progenitor death and disrupted neuron numbers) in our Scz organoids, our next goal was to identify and define the molecular candidates responsible for these effects. To do this, we employed high-throughput, unbiased, and computational tools to deconvolute the molecular architecture of Scz organoids. A major goal of this approach was to resolve novel Scz disease signatures at the posttranslational level. Additionally, we also sought to determine whether known disease factors and pathways from the Scz GWAS literature were recapitulated in Scz patient-derived organoids.

To map the proteome of 3D human-derived organoids, we adapted Tandem Mass Tag (TMT) chemistry to globally barcode peptides isolated from Ctrl and Scz cerebral organoids. This allowed a pseudorandomly selected cross-section of organoid samples to be pooled into a single multiplexed suspension that could be simultaneously subjected to quantitative Liquid Chromatography-Mass Spectrometry (LC-MS; fig. 2a). This approach identified 3772 proteins, of which ∼97.4% were quantifiable. The most abundantly expressed proteins, evidence of neuronal enrichment, and differentially expressed candidates are presented in Supplementary Tables 2-7. Remarkably, the overlap in protein identities between Ctrl and Scz cerebral organoids was almost complete (99.95% similarity in protein diversity) with only two proteins being unique to Scz cerebral organoids (WASL & MPV17; fig. 2b and fig. S3 for further data and discussion). The reproducibility of samples was robust in this dataset (fig. S3d), and even at the single-protein level coefficients of variation remained low for over 3,000 identified peptides (fig. 2c). Indeed, at baseline, only 3 of 3671 (or, ∼0.08% of the entire organoid proteome) exhibited substantial variability. Therefore, while undirected cerebral organoids varied in their morphologies and appearance, their macromolecular composition was found to be relatively stable and consistent. Additionally, both Ctrl and Scz cerebral organoids exhibited enrichment for neuronal and brain developmental pathways (e.g., axons, neurogenesis, postsynaptic density, neuron projection development and WNT signaling; see Supplementary Table 3). This once more provided independent confirmation that neural induction was universally achieved in all iPSC lines.

**Figure 2.**
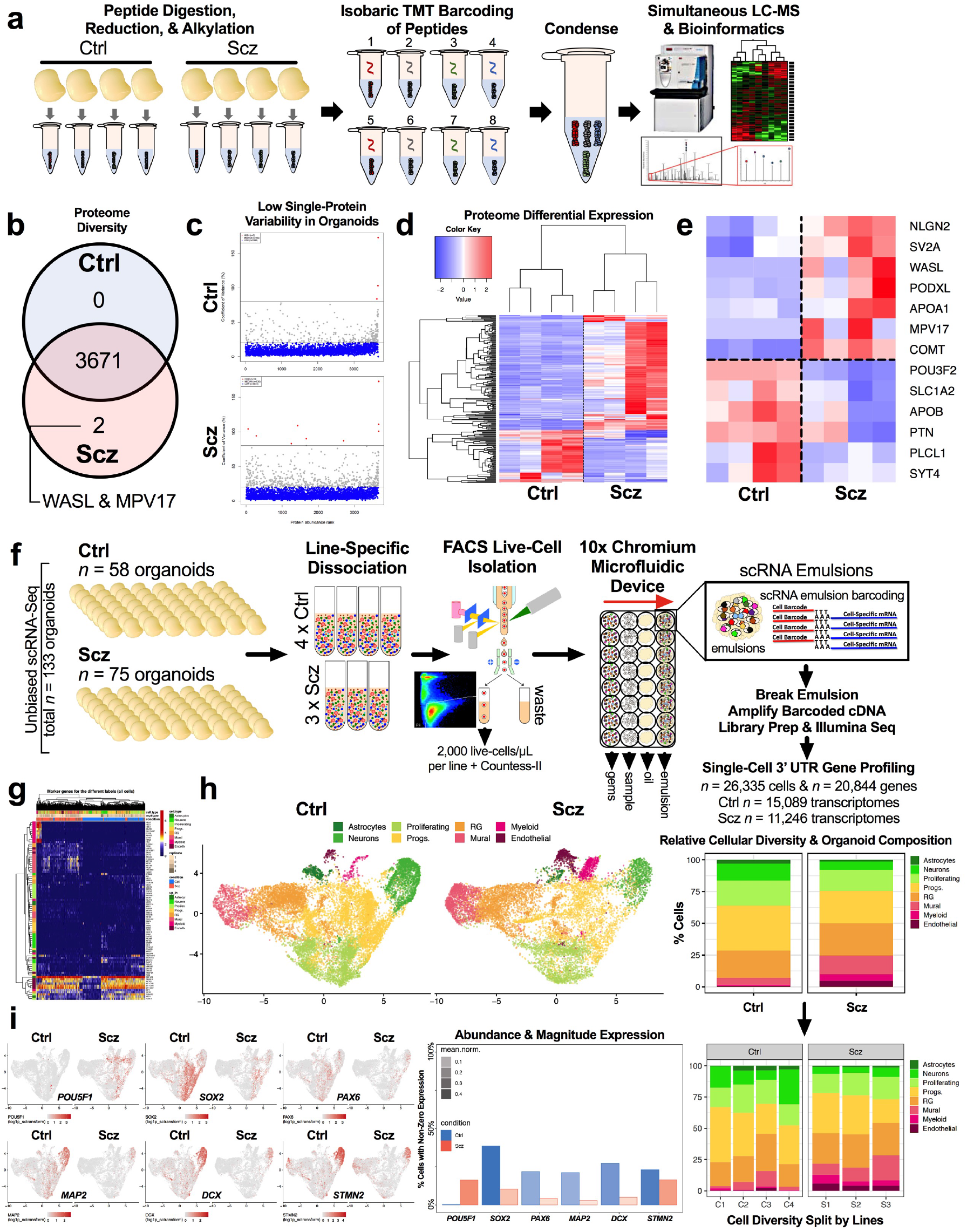
Proteomics and single-cell transcriptomics identify novel disease signatures of Scz patient organoids. a, Schematic of proteome mapping pipeline using Tandem Mass Tag (TMT) chemistry and Liquid Chromatography-Mass Spectrometry (LC-MS). Briefly, whole organoids were subjected to tryptic digestion to isolate proteins from each sample. Proteins were next isobarically barcoded with TMT reagents, and then condensed into a single multiplexed suspension. This allowed samples to be run concurrently, eliminating batch-specific technical variance, and a low-noise assessment of both organoid variability (using a 4x2 design) and the identification of translated disease factors. Multiplexed suspensions were subsequently subjected to quantitative TMT-LC-MS for peptide detection, deconvolution, and quantification, and then computationally analyzed. **b**, **Scz organoids recapitulate the developmental proteome.** Given numerous cellular signatures of Scz that separated cases from Ctrls (fig. 1), we sought to determine the proteome diversity of Ctrl and Scz 3D organoids. The Scz organoid proteome was 99.95% identical in pooled protein diversity relative to Ctrls. Thus, Scz proteome differences are likely to be explained not in the induction of differential factors at the posttranslational level, but rather their total molecular quantity. **c, Individual proteins exhibit low variability in expression between groups**. Given the high degree of conservation in peptide diversity between both Ctrl and Scz organoids, we next examined dispersion of individual proteins by generating their coefficient of variation (CVs) for each group. In both Ctrl and Scz samples, over 3000 detectable peptides for each group exhibited low variability (<20% CVs; blue dots). This reflects that Ctrl and Scz organoids exhibited similar patterns of proteome reproducibility. **d, Proteome-wide clustering separates Scz patient-derived organoids from Ctrls.** A proteome-wide heat-map shows differences in LC-MS expression intensities between groups, which unveils clustering of differentially expressed proteins in Ctrl versus Scz organoids. Specifically, Scz organoids are defined by the differential expression of just 222 peptides, or ∼5.9% of the detected proteome. Thus, consistent with our hypothesis, Scz organoid samples principally differed in their quantity, rather than diversity, of developmental factors. **e, Disease factors are detected and differentially expressed in Scz organoids.** A heat-map of selected differentially expressed proteins in Scz organoids (Log2 key from panel f still applies). This included POU-domain peptide fragments that putatively mapped to the forebrain neuronal-development factor POU3F2 (BRN2), and disease factors with known and/or likely involvement in disease pathophysiology (e.g. COMT/PLCL1). Additionally, factors with putative genetic risk but otherwise unestablished disease biology (e.g. PTN) were also differentially expressed in Scz organoids. **f, Schematic of pipeline for live single-cell RNA sequencing of organoids.** Briefly, organoids from 4 Ctrl and 3 Scz lines were generated concurrently, pooled by- line, and dissociated to a single-cell suspension. We rapidly conducted survival-based high-throughput FACS to purify samples to 2000 live cells/µm per line. For robustness, post-FACS live cell viability was also cross-confirmed using Countess-II. Live cell suspensions were next rapidly loaded into 10x chromium microfluidic devices to produce barcoded single-cell nanodroplet emulsions. This emulsion was later broken, barcoded samples were amplified, libraries prepared, subjected to Illumina sequencing, and then parsed through a suite of unbiased computational analyses. **g, Clustering of 26,335 transcriptomes recapitulates fetal brain cellular identities.** Heatmap of marker genes of individual clusters identified via pairwise comparisons of the normalized expression values for cells of a given cluster vs. the cells of all other clusters. Shown here are unbiased gene sets, which define the top 10 gene markers for each cluster that met a high-pass FDR threshold of 1% and >15,000 total read counts. Many of these prototypic markers defined cell-types consistent with human fetal tissue [1, 2] (see below). **h, Resolving the identities of differentially enriched cell-types in Scz organoids.** Shown are UMAP coordinates for 26,335 transcriptomes split by Ctrl and Scz cases. Cell-type labels were determined via a variety of approaches including marker gene- expression, automated annotation, and, namely, comparison with human fetal samples [2]. Bar chart (right) depicts cell-type proportions, illustrating that ∼93% of Ctrl scRNA- Seq transcriptomes were identified as neural progenitors, proliferating cells, or terminal cortical cell-types (e.g. neurons and glia). Compared to this, only ∼75% of Scz cell-types exhibited a similar conservation of identity. Compared to remaining cell types in Ctrl organoids (∼7% cells), the remaining ∼25% of Scz scRNA-Seq transcriptomes reflected enrichment for brain-related cell-types including putative neuro-endothelial cells, structural markers, developing vasculature, retinal, and choroid plexus markers in Scz organoids. This analysis therefore revealed the cell-types produced at the expense of neurons in Scz organoids alluded to in pulse-chase experimentation depicted in fig. 1e. Of note, all cell-types exhibited reproducible proportions across individual iPSC donors within respective groups (i.e., all Scz organoids exhibited similarly reproducible alterations in cell-type diversity, which was defined by an overarching loss of neurons). **i, Confirmation of progenitor and neuronal depletion in Scz organoids.** Scz organoids exhibited a striking enrichment of the pluripotent stem cell marker *POU5F1* (OCT4), which overlapped *SOX2+* and *PAX6*+ progenitor UMAP coordinates. Correspondingly, Scz organoids exhibited depletion of progenitor markers (*SOX2*+ and *PAX6*+) as well as the pan-neuronal markers (*MAP2*+, *DCX*+, and *STMN2*+). These expression differences reflect both abundance and magnitude. Thus, Scz organoids exhibit a deleterious increase in pluripotent cell-types (see *POU5F1*/OCT4) in progenitor clusters, but an overall depletion of progenitors and neurons. For fig. 2f-j, total *n* = 26, 335 transcriptomes, *n* = 20,844 genes from 7 iPSC lines; Ctrl *n* = 15,089 transcriptomes from 4 Ctrl iPSC lines, and Scz *n* = 11,246 transcriptomes from 3 Scz iPSC lines. Ctrl: Control, Scz: Schizophrenia.

Although the diversity of proteins in Ctrl and Scz organoids were very similar, further computational analysis revealed notable differences in the quantity of peptides within Scz organoids. Analysis of global Log2 intensities revealed that 222 proteins, reflecting ∼5.9% of the proteome, differed in expression between Ctrl and Scz organoids (fig. 2d). Of these proteins, 151 were up-regulated and 71 were down-regulated. Interestingly, peptide fragments of POU-domain containing transcription factors were identified in Scz organoids. This is of note, as the upper-layer cortical neuron marker and forebrain-specific transcription factor, POU3F2 (BRN2), is prominently involved in embryonic corticogenesis (see identity markers in [38]). BRN2 is an established initiator of neuronal differentiation [71] and BRN2 is also dysregulated in organoids with DISC1 disruption [72]. When differentially expressed proteins were screened against large-scale Scz gene association studies [36, 73], at least ∼70 putative Scz risk proteins were discovered to be commonly present in organoids. Only 4 of these specific risk factors, PLCL1, PTN, PODXL and COMT, displayed significant differential expression in Scz organoids.

Notably, PLCL1 (phospholipase-like C1; involved in GABAergic neurotransmission [74]) and PTN (pleiotrophin; a possible neurodevelopmental growth factor [75]) were significantly down-regulated in Scz organoids. Contrary to this, PODXL (podocalyxin; a neural adhesion molecule involved in axonal fasciculation, neurite outgrowth, and synaptogenesis [76]) and COMT (catechol-O-methyltransferase; involved in dopamine and catecholamine elimination [77] and cognitive response to antipsychotics [78]) were innately up-regulated in Scz organoids (fig. 2e). This suggests that neuropathology of Scz converges upon the disruption of common molecular factors with established disease risk, perhaps as a consequence of a shared disruption of broader regulatory networks.

### Single-cell sequencing reveals aberrant cell-type diversity in Scz organoids

Next, to study disease signatures of Scz with greater resolution, both with respect to cell-specificity and capture potential, we employed scRNA-Seq. To do this, we used chromium microfluidic devices to generate single-cell nanodroplet emulsions that could then be broken and amplified for library preparation and sequencing (fig. 2f).

Our high-content scRNA-Seq pipeline yielded 26,335 single-cell transcriptomes, comprising 20,844 captured genes that could be harmonized into 8 cellular clusters (fig. 2g-h) identified against human fetal brain samples [81]. Replicating data from fig. 1-2, there was decreased abundance of both progenitors (*SOX2* and *PAX6*) and neurons (*MAP2, DCX*, and *STMN2*) in Scz organoids (fig. 2h-i). Instead, Scz organoids exhibited abnormal enrichment for brain-related cell-types including neuroendothelia, myeloid, mural, choroid plexus, and retinal markers (fig. 2h). Ctrl samples also exhibited these same cell-types but in lower abundance (∼7% of all Ctrl cell-types, vs. ∼25% of Scz cell-types; fig. 2h). Scz organoids therefore recapitulated all appropriate cell-types also present in Ctrl organoids but, due to disrupted differentiation (fig. 2e), Scz organoids were defined by an increase in neuroendothelial-related cells and a decrease in neuron numbers.

### Altered pseudotime trajectories reveals remodeling of Scz organoid cell-lineages

We next sought to determine whether alterations in differentiation and diversity in Scz organoids arose from transcriptomic remodeling of cell lineages. We computationally modeled the differentiation trajectories of organoids by harmonizing single-cell transcriptomes into principal components that exist in pseudotime [84]. This yielded differentiation trajectories of Ctrl and Scz organoids (fig. 3a) that were similar but subtly distinct (fig. 3b). Ctrl organoids reflected enrichment of known neuronal programming factors with established mechanistic roles in neuronal differentiation (fig. 3b-c). Contrary to this, Scz organoids were defined by different markers that were instead associated with pluripotency, cytoskeletal filaments, and neuroendothelia (fig. 3b). This included *POU5F1* (*OCT4*), the proliferation and apoptosis balancing factor *CRABP1,* and pan enrichment for the inflammatory factor *IFITM3* in Scz organoids (fig. 3c vs. 3a for overlap). In Scz organoids, the enrichment of *POU5F1*+ cells coincided with cell-type clusters that were broadly consistent with progenitors and developing neurons (see overlap of markers in fig. 3c vs. clusters in 3a), indicating that these markers were expressed within cell-type clusters that exhibited evidence of neural induction. This indicates that these aberrant markers were over-populated in progenitor and neuronal clusters at the expense of other prototypic neuronal development factors and related cell-types (see UMAPs in fig. 2i). This analysis was also consistent with other up- and down-regulated developmental factors in Scz organoids identified during unbiased clustering and analysis (see fig. 2i and S4). Therefore, this statistical pseudotime analysis provided unbiased confirmation that Scz cell-lineages exhibited remodeling that diverts progenitor differentiation away from neuron production within organoids.

**Figure 3.**
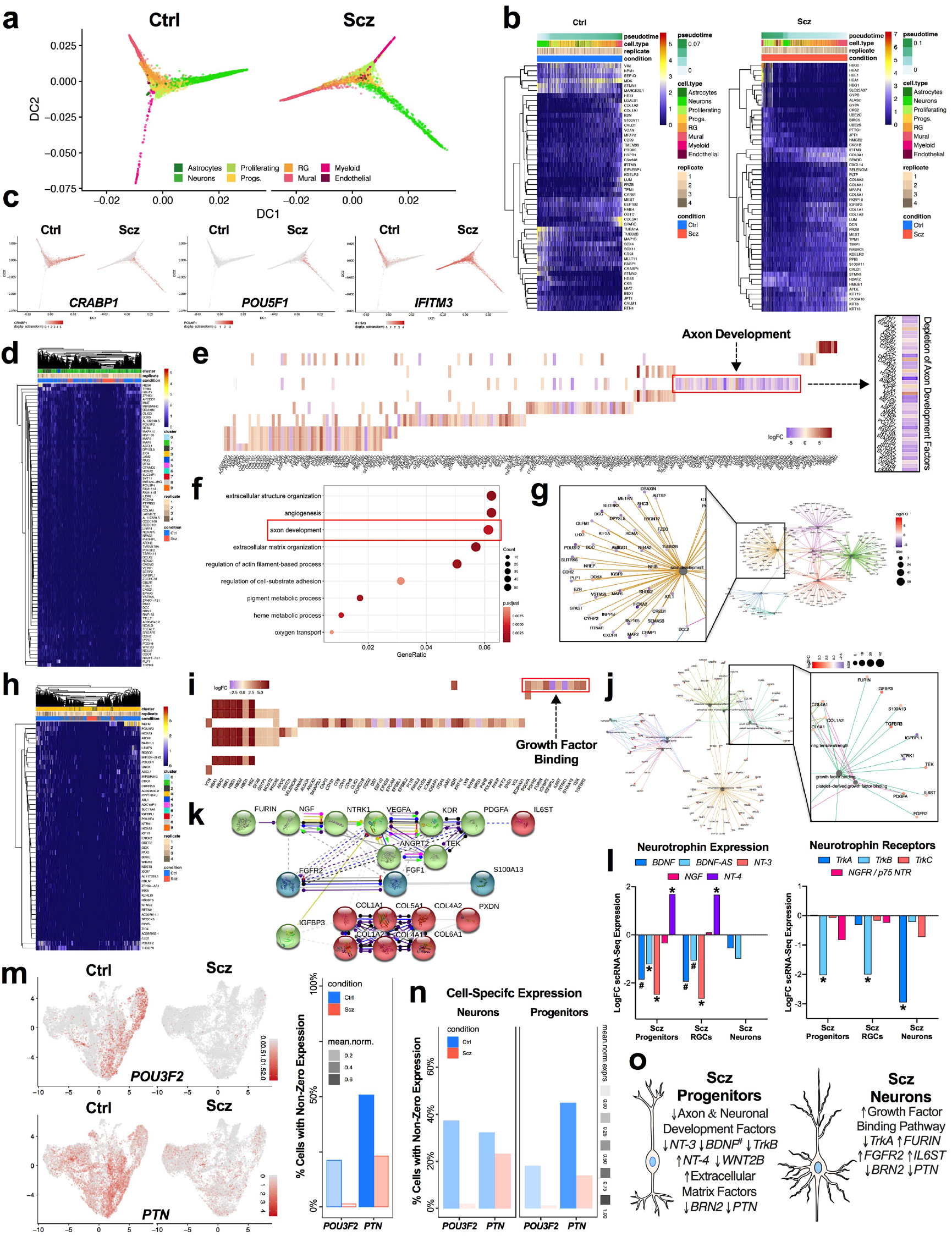
Altered cell-lineages and cell-specific neuropathology in Scz organoids. a-c, Computational analyses reveal altered single-cell lineages in Scz organoids. Diffusion maps of single-cell differentiation trajectories were generated for all 26,335 cells via harmonized principal components. This revealed similar, but subtly altered, differentiation trajectories in Scz organoids (a). Heatmaps of slingshot-identified genes most-associated with pseudotime cell lineages of Scz organoids (b). Note differences in enrichment for collagen and matrix organization factors in Scz single-cell transcriptomes, whereas Ctrl organoids tended to exhibit up-regulation of neuronal factors (e.g. *STMN2*, *TUBB2B*, *CRABP1*) as expected. Expression patterns of selected genes of interest, based on statistical thresholds, are projected onto diffusion map coordinates (c). Notably, markers such as *CRABP1, IFITM3,* and *POU5F1* (OCT4) segregated Scz from Ctrl transcriptomes within differentiation trajectories (c), and were inappropriately enriched in progenitor cell-types and neuronal trajectories (see reference cell-type clusters in panel a). Therefore, these markers did not identify a broader pool of undifferentiated stem cells, but rather indicated a series of intrinsic alterations within pseudotime trajectories of Scz cell-types and their differentiation patterns. d-g, Scz progenitors exhibit disrupted transcription of axon development factors. *SOX2*+ progenitor scRNA-Seq transcriptomes were isolated and differentially expressed genes (DEG) examined. Heatmap denotes unbiased-discovery of downregulated DEG targets in Scz progenitors (d; 5% FDR threshold average logFC Scz/Ctrl ≤ 2.5, and >3 for up-regulated analysis in fig. S4d). This revealed down-regulation of a gene-set involved in axon development (e; see inset). Other DEG targets in Scz progenitors by biological process are presented here as a dotplot (f), resulted in gene-set enrichment for extracellular organization, adhesion, as well as angiogenesis in Scz progenitors. Gene ontology biological processes are shown in spoke-plot (g), and axon factors are enlarged in isobar-linked image for emphasis and evaluation (inset of e and g). **h-k, Scz neurons exhibit pathway enrichment for growth factor binding.** Similar to progenitors, neuron scRNA-Seq transcriptomes were examined for DEGs. Heatmap denotes unbiased-discovery of downregulated DEG targets in Scz neurons (h; 5% FDR threshold average LogFC Scz/Ctrl ≤ 2.5, and >3 for up-regulated analysis depicted in fig. S4e). Analysis of gene ontologies by molecular function revealed enrichment for growth factor binding in Scz neurons, which was defined by downregulated TrkA (*NTRK1*) neurotrophin receptor gene expression and upregulation of factors such as FURIN (neurotrophin pro→mature processing), *FGFR2*, *PDGFA*, *TGFBR3,* and the interleukin-6 signal transducer *IL6ST* (i-j). Protein-protein interactions between enriched growth factor binding factors in Scz neurons revealed notable interplay and interaction pathways (e.g. Furin→NGF→NTRK1) that place neuron-specific DEGs such as IL6ST and S100A13 as downstream targets of growth factor binding enrichment (k). Alternatively, these downstream factors may be putatively repressed by the inhibitory interaction effect of IGFBP3 on VEGFA (k), which are prototypic markers of neurovascular cells which were also upregulated in their abundance within Scz organoids (see fig. 3h). Other pathway analysis (e.g. KEGG) identified altered Pl3K-Akt signaling in Scz neurons (see fig. S4g). **l, Cell-specific disruption of neurotrophic factors and receptors in Scz organoids.** Given unbiased detection of alterations in *NT-3* (*NTF3*) in Scz progenitors and *TrkA* (*NTRK1*) receptors in Scz neurons, we next examined progenitor cell-types for specific differences in neurotrophins (*BDNF*, *BDNF-Antisense*/*BDNF-AS*, *NGF*, *NT-3*, and *NT-4*; left) and their cognate tyrosine kinase receptors (*TrkA*, *TrkB*, *TrkC*, & *p75 NTR/NGFR*; right). *SOX2*+ progenitors and radial glial cell (RGC) progenitors examined almost identical neurotrophin dysregulation and were defined by the trend for decreased expression of *BDNF*, *BDNF-AS, NT-3,* and *TrkB* expression. Curiously, *NT-4* was upregulated in Scz progenitor cell-types. Scz neurons were defined only by their near 3- fold reduction of *TrkA* gene expression. m-n, Validating gene-expression differences of candidate proteome-derived targets, *POU3F2* (BRN2) and *PTN*, in Scz organoids. Candidate proteome targets (fig. 2e) were cross-examined for differences at the single-cell level (see fig. S5 for expression of proteome targets in global, progenitor-only and neuron-only scRNA-Seq datasets). Given survival, differentiation, and diminished growth factor support, here we emphasize the neuronal transcription factor BRN2 (*POU3F2*) and putative growth factor PTN as putative targets for mechanism experiments. Scz progenitors exhibited almost complete depletion of BRN2 relative to Ctrl progenitors, indicating disrupted induction of neuronal differentiation. PTN expression was disrupted in both Scz progenitors and neurons but exhibited a greater difference in progenitors. o, Resolving neural cell-type specificity of *POU3F2* (BRN2) and *PTN* depletion in Scz organoids. Schematic summary of cell-specific alterations in Scz progenitors (left) and neurons (right). Scz progenitors exhibited entropy in neurotrophic growth factors, specific reduction of TrkB (*NTRK2*), and depletion of axon development factors. Scz progenitors also exhibit enrichment for extracellular structure, matrix organization, and angiogenesis factors, which partially explains the intrinsic alteration in cell lineage (fig. 3a-c) and differentiation trajectories towards neuro-endothelial factors and cell types (fig. 2 and 3). Total scRNA-Seq dataset comprised *n* = 26, 335 cells, *n* = 20,844 genes from 7 iPSC lines; Ctrl *n* = 15,089 scRNA-Seq transcriptomes from 4 Ctrl iPSC lines, and Scz *n* = 11, 246 scRNA-seq transcriptomes from 3 Scz iPSC lines. * significant *p* and FDR values. # = *p* value significant, with a final FDR score at the cut-off threshold. Ctrl: Control, Scz: Schizophrenia.

### Cell-specific encoding of neuropathology in Scz progenitors and neurons

We next sought to define the specific transcriptional factors responsible for the disrupted neurogenic potential of *SOX2*+ Scz progenitors (fig. 3d-g and S4d). Numerous transcriptional regulators of differentiation were down-regulated in Scz progenitors (fig. 3d), including a depletion of axon development pathway factors (fig. 3e-f). Intriguingly, delayed axon development is also a known sequelae of *in utero* animal models of Scz [85]. Many of the axon development factors identified here also regulate neuronal differentiation and development (e.g. *HOXA2*, *AUTS2*, *MAP2*, *SLITRK6*, *WNT2B, NRN1;* see inset of fig. 3g). Prominently, this included the depletion of *POU3F2* (BRN2) in Scz progenitors. Down-regulation of Neurotrophin-3 mRNA (*NT-3* or *NTF3)* was also detected in Scz progenitors. Consistent with pseudotime analyses, Scz progenitors exhibited abnormal enrichment for structural, filament, adhesion, and angiogenic pathways (see fig. 3f). Therefore, we identify that Scz progenitors exhibit depletion of an ensemble of development factors that promote neuronal differentiation and potentiate early neuronal development.

We next sought to determine if developing neurons also exhibit cell-specific signatures of Scz. Similar to progenitors, several neuronal development factors remained down- regulated in Scz neurons including *POU3F2* (BRN2). Neuronal filament factors (e.g. *NEFM*) and guidance factors (e.g. *ROBO3* and *NTNG2*; see fig. 3h) were also disrupted in Scz neurons. Curiously, we identified enrichment for Pl3K-Akt signaling (see fig. S4f) as well as a growth factor binding pathway in Scz neurons (fig. 3i-j). This included down- regulation of the *TrkA* (*NTRK1*) receptor and the up-regulation of the neurotrophin cleavage factor, *FURIN*. In humans, *FURIN* localizes to one of 108 Scz loci with genome-wide significance [36]. In support of this, interactome modeling revealed potential interplay between growth factor binding factors that may be initiated by interactions related to FURIN and TrkA at the posttranslational level (fig. 3k). The interleukin-6 signal transducer (*IL6ST*), which was up-regulated in Scz neurons, may be downstream of *FURIN* and *TrkA* based on interactome modeling (fig. 3k). Notably, FURIN is responsible for cleaving immature neurotrophins into their mature isoforms, and the differential expression of these identified pathways (e.g. growth factor binding and Pl3K-Akt signaling factors) is broadly consistent with a compensatory response to diminished neurotrophin signaling. This suggests that growth factor binding factors, including *FURIN*, may be enriched in neurons to compensate for broadly diminished growth factor signaling via *TrkA*, which is a known regulator of neuronal differentiation, development, and survival.

### Cell-specific neurotrophin alterations in Scz progenitors and neurons

The down-regulation of *NT-3* in Scz progenitors and *TrkA* in Scz neurons in unbiased statistical modeling indicates that other discrete patterns of neurotrophin dysfunction may exist. Deeper hypothesis-driven analysis revealed that Scz progenitors also tended to exhibit down-regulated *BDNF*, but up-regulated *NT-4,* gene-expression.

Correspondingly, we identified that *TrkB (NTRK2*) was the only down-regulated neurotrophin receptor in progenitors (fig. 3l). Contrary to this, Scz neurons only exhibited a cell-specific down-regulation of *TrkA* (as described above; fig. 3l). These data thus provide further evidence to support our hypothesis that divergent cell-specific neuropathology may converge upon common pathways in Scz organoids (e.g. growth factor support), resulting in related but non-overlapping cell-specific neuropathology of Scz between progenitors and neurons.

### Identifying BRN2 and PTN as candidate disease factors in organoids

To identify potential mechanistic factors of Scz neuropathology in our patient-derived organoids, we examined whether proteome targets could be validated in our scRNA-seq transcriptomic datasets (see fig. S5a-b). Global, as well as progenitor- and neuron-specific, analyses revealed that *POU3F2* (*BRN2*) and *PTN* were depleted in Scz samples (fig. 3m-n). Intriguingly, *PTN* holds disease relevance due to the proximity of an index SNP/novel locus with genome-wide significance in the largest Scz GWAS conducted to date (*n* = ∼150,000 humans, see [36] and fig. S6). On the other hand, BRN2 does not appear to be a genetic locus of Scz risk, but does exhibit some evidence of disease linkage in prior differential methylation studies (fig. S6c). This suggests that BRN2 may be a commonly targeted downstream pathophysiological factor (see Scz database repository mining in fig. S6). The neurobiology of these candidates only further amplified our interest, as BRN2 promotes the production of late-born neurons during embryonic development [71], and PTN is a putative growth factor that promotes cell survival [92] as well as other neurodevelopmental processes. Importantly, informatic mining of the BrainCloud^TM^ (dbGaP Accession: <u>phs000417.v2.p1</u>) database revealed that both BRN2 and PTN exhibit peak expression during prenatal human brain development (*n* = 269 time-points including *n* = 38 fetal samples, see fig S6a), making them bonafide embryonic targets that may be related to Scz risk during this period. In sum, these supplementary informatics analyses provided additional veracity for our core Scz phenotypes, providing confidence in the translational value of our 3D organoid cultures and their ability to recapitulate disruption of novel embryonic disease factors.

### Modulating BRN2 levels in Scz progenitors rescues neurogenesis

Having established the reproducibility of our core phenotypes (progenitor death and neuronal loss) in Scz organoids, we next sought to mechanistically establish the functionality of the candidate rescue factor BRN2 within our Scz organoid system. BRN2 is a forebrain specific transcription factor expressed in neural progenitors as well as in migrating neurons in the mammalian brain (see fig. 3 and [71]). When disrupted *in vivo,* stalled production of upper-layer neurons ensues, indicating a functional role for BRN2 in early cortical neurogenesis [71]. The endogenous disruption of BRN2 in Scz organoids (fig. 2-3) implicated that BRN2 may be a downstream pathophysiological substrate of Scz neuropathology. We hypothesized that the robust down-regulation of BRN2 in Scz progenitors resulted in the disrupted neuronal differentiation observed in Scz organoids. To test this, we specifically supplemented BRN2 in Scz progenitors and assessed neuron numbers within the same Scz organoids. To do this, we used a sophisticated self-regulating viral construct (hPGK>BRN2:P2A:EGFP.MIRT124^4^, see fig. S7a-c; hitherto *BRN2*-Virus) that maintained sustained production of exogenous BRN2-GFP expression within infected progenitor cells but not post-mitotic neurons (fig. 4a-b, S7). This was achieved via a neuronal “off-switch” in viral transcripts, which was controlled by four recognition sites for the neuronal noncoding RNA miRNA-124 (MIRT-124^4^; for details see Methods). During the early stages of organoid development, tissue was infected with either a control-GFP (*CTRL*-Virus) or *BRN2*-GFP lentivirus (fig. 4a), leading to viral expression within neural progenitors as expected (fig. 4b). Whole-organoid images revealed that Scz phenotypes, such as neuronal loss, were independently recapitulated in Scz organoids (fig. 4c, see also fig. S8 for enlarged images of neuronal loss). Analysis revealed that *BRN2*-Virus+ Scz organoids exhibited a significant increase in BRN2+ late-born neuron numbers. Indeed, late-born neuron numbers were rescued to levels that were consistent with Ctrl organoids (fig. 4d). In addition, MAP2+ neuron numbers were also increased in *BRN2*-Virus+ Scz organoids relative to *CTRL*-Virus+ Scz organoids (fig. 4e). However, viral delivery of exogenous BRN2 to Scz progenitors failed to rescue increased rates of progenitor apoptosis within Scz organoids (fig. 4f).

**Figure 4.**
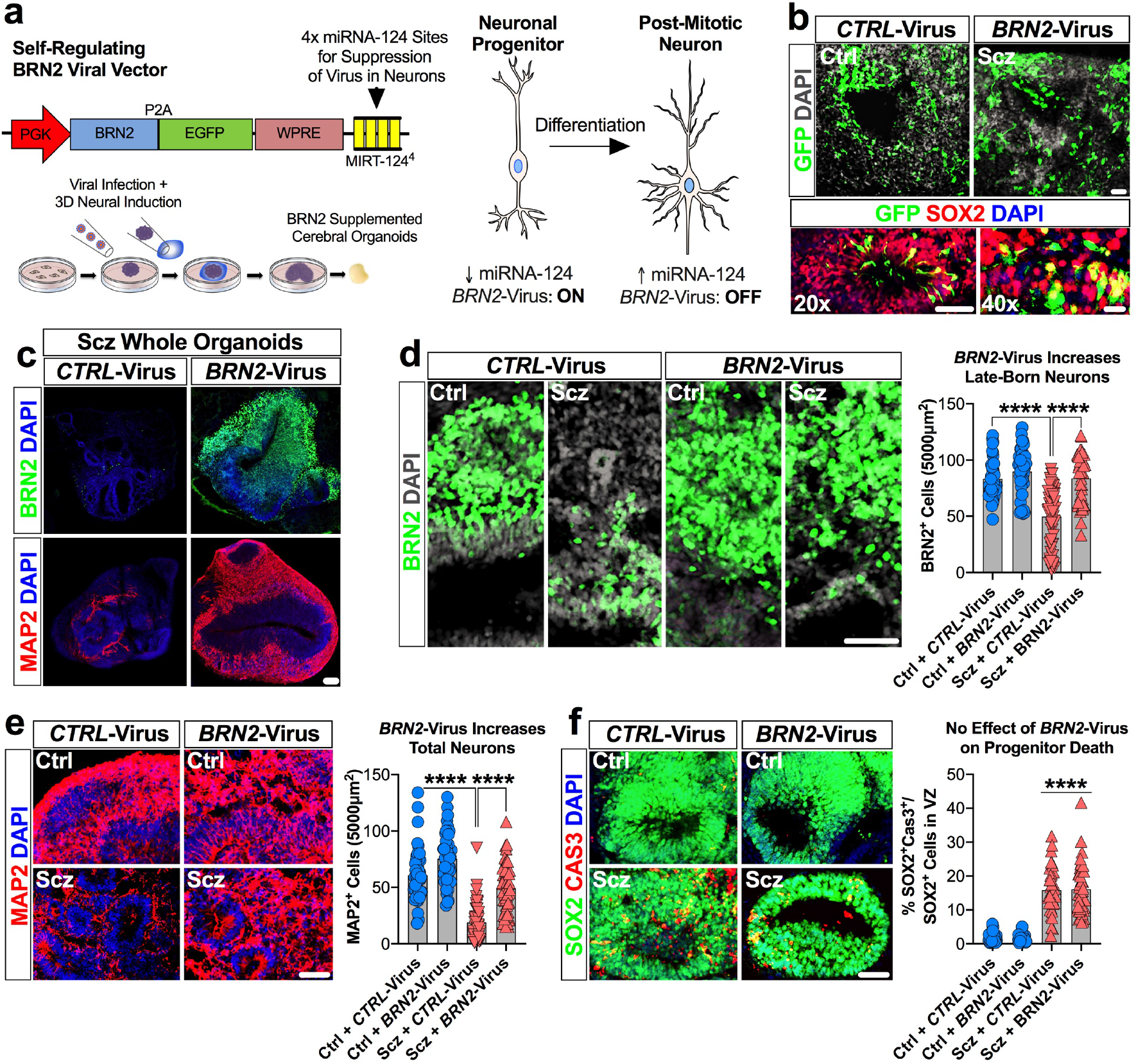
BRN2 rescues neurogenesis, but not survival, in Scz organoids. a, Schematic of self-regulating *BRN2* lentiviral vector and supplementation strategy for BRN2 rescue experiments. Briefly, we modified a previously validated lentiviral construct [7] that transiently induces exogenous BRN2 expression but “switches-off” upon completion of neuronal differentiation and assumption of a post-mitotic neuronal fate [7]. After supplementing neuronal transcriptional programs, exogenous *BRN2*-Virus transcripts are decayed via binding of the neuron-specific noncoding RNA, miRNA-124, to recognition units embedded within viral transcripts (see fig. S7). This *BRN2*-Virus construct thus allows transient supplementation of BRN2 levels in Scz progenitors undergoing differentiation without sustained overexpression of this target in mature neurons. **b-e, BRN2 supplementation increases neuron numbers in Scz organoids.** Ctrl and Scz organoids were infected with *CTRL*-Virus and *BRN2*-Virus during organoid neural induction. Images show GFP+ cells within ventricular zones. Viral infection rates appeared similar between Ctrl and Scz organoids, and viral GFP expression was detected within progenitor pools as expected (b). Scz organoids infected with *CTRL*- Virus exhibited fewer neurons than Ctrl organoids. However, when comparing Scz organoids infected with *CTRL-*Virus versus *BRN2-*Virus, we observed a significant rescue of BRN2+ neuron number. Scz organoids infected with *BRN2*-Virus exhibited substantially increased BRN2+ neuron numbers, which were comparable to Ctrl organoids. Enlarged whole-organoid images are provided in Supplementary Material (see fig. S8), and graphs reflect raw data for complete data transparency of rescue effects (d, *CTRL*-Virus Ctrl organoids *n* = 43 fields, *n* = 16 organoids, and *n* = 3 independent Ctrl lines; *BRN2*-Virus Ctrl organoids *n* = 40 fields, *n* = 18 organoids, and *n* = 3 Ctrl independent lines; *CTRL*-Virus Scz organoids *n* = 65 fields, *n* = 25 organoids, and *n* = 5 independent Scz lines; *BRN2*-Virus Scz organoids *n* = 42 fields, *n* = 14 organoids, and *n* = 4 independent Scz lines). To determine if this effect was reflected in pan neuronal numbers, we also examined MAP2+ neurons in infected organoids. Consistent with data in fig, 2, *CTRL*-Virus+ Scz organoids exhibited a decreased number of MAP2+ neurons compared to Ctrl organoids. However, MAP2+ neuron numbers were significantly increased in the *BRN2*-Virus+ Scz organoids (e, *CTRL*-Virus Ctrl organoids *n* = 73 fields, *n* = 35 organoids, and *n* = 5 independent Ctrl lines; *BRN2*-Virus Ctrl organoids *n* = 55 fields, *n* = 26 organoids, and *n* = 3 Ctrl independent lines; *CTRL*-Virus Scz organoids *n* = 52 fields, *n* = 25 organoids, and *n* = 5 independent Scz lines; *BRN2*- Virus Scz organoids *n* = 64 fields, *n* = 28 organoids, and *n* = 5 independent Scz lines). Thus, transient BRN2 supplementation resulted in a significant recovery of neurons in 3D Scz patient-derived organoids, which confirms a mechanistic role for BRN2 within Scz organoids. Each data point on graphs reflects raw data (an independent, non- overlapping, cortical field) for complete data transparency, with the average of individual iPSC lines provided in Supplementary Material (see fig. S7d) **f, No effect of BRN2 supplementation on progenitor cell death in Scz organoids.** Scz organoids are associated with increased rates of cell death of ventricular zone neural progenitors (fig. 1). To determine if BRN2 regulates the survival of progenitors, we assessed the number of CAS3+ in infected organoids. In both *CTRL-* and *BRN2*-Virus infected Scz organoids, there was an increase in progenitor death relative to Ctrl samples (e, *CTRL*-Virus Ctrl organoids *n* = 34 fields, *n* = 19 organoids, and *n* = 3 independent Ctrl lines; *BRN2*-Virus Ctrl organoids *n* = 35 fields, *n* = 19 organoids, and *n* = 3 independent Ctrl lines; *CTRL*-Virus Scz organoids *n* = 35 fields, *n* = 20 organoids, and *n* = 4 independent Scz lines; *BRN2*-Virus Scz organoids *n* = 45 fields, *n* = 23 organoids, and *n* = 4 independent Scz lines). These data indicate that decreased levels of BRN2 do not contribute to the increased apoptosis of progenitors in Scz organoids. Each data point on graphs reflects raw data (comprising an independent ventricular zone) for complete data transparency (for the average of groups, see fig. S7d). **h, Schematic of BRN2 rescue experiments in Scz cerebral organoids.** Here we find that BRN2 has a mechanistic role in promoting neuron production in Scz organoids, but not the survival of neuronal progenitors or neurons themselves. Correspondingly, cells with red nuclei represent apoptotic cells in schematic. This selective rescuing effect of BRN2 highlights that multiple factors and pathways likely combine to produce progenitor and neuronal pathology in developing cortical assemblies of Scz organoids \*\*\*\**p* < 0.0001. Error bars reflect Standard Error of the Mean. Scale bar: b-20x = 60µm, b-40x = 20µm, c-f = 60µm. Ctrl: Control, Scz: Schizophrenia. X in schematic denotes cell death.

While raw data are presented (fig. 4) for transparency, further analysis of phenotypes by the average of individual iPSC lines reaffirmed that *BRN2*-Virus Scz organoids exhibited increased neuron numbers but no change in progenitor death (see fig. S7). Together, these data establish that down-regulated BRN2 mechanistically contributes to neuronal differentiation, but not apoptosis, phenotypes in Scz organoids (fig. 4h). Therefore, due to our inability to rescue cell death phenotypes with *BRN2*-Virus, we considered the possibility that other disease mechanisms must be operating within Scz organoids.

### PTN corrects neuron numbers in Scz organoids via neurotrophic-like effects

Recent work has established that PTN exhibits strong expression in neural progenitors (see [90] and fig. S5a). We were also able to independently replicate the expression of PTN within progenitors here via scRNA-Seq (fig. 3n). PTN has been suggested to play a role in early corticogenesis by promoting neuronal differentiation [91]. PTN can also protect neurons from apoptosis *in vivo* [92]. To determine whether PTN plays a role in early Scz pathology, we adapted recombinant human PTN as a morphogen to generate Vehicle- and PTN-Supplemented Ctrl and Scz cerebral organoids (fig. 5a). Similar to earlier experiments (fig. 1, 2, 4), Scz phenotypes were recapitulated in Vehicle-treated Scz organoids and this included a significant increase in Scz progenitor apoptosis (fig. 5c). However, relative to this, PTN-treated Scz organoids exhibited a significant reduction in progenitor apoptosis (fig. 5c, and S9a for enlarged whole-organoid images). PTN-treated Scz organoids exhibited increased new-born cell survival (fig. S10) and thus significantly increased neuronal differentiation (fig. 5d). This was reflected in the numbers of MAP2+ neurons, which were increased relative to Vehicle-treated Scz organoids (fig. 5e; see also enlarged whole-organoid images in fig. S9b). Importantly, these phenotypes were replicated irrespective of whether data were analyzed as a pool or by the average of individual iPSC lines (see fig. 5 and S11). In sum, these experiments collectively established that PTN mechanistically regulated new-born cell survival and thus neuron number in Scz organoids by simultaneously promoting neurogenesis and restraining apoptosis.

**Figure 5.**
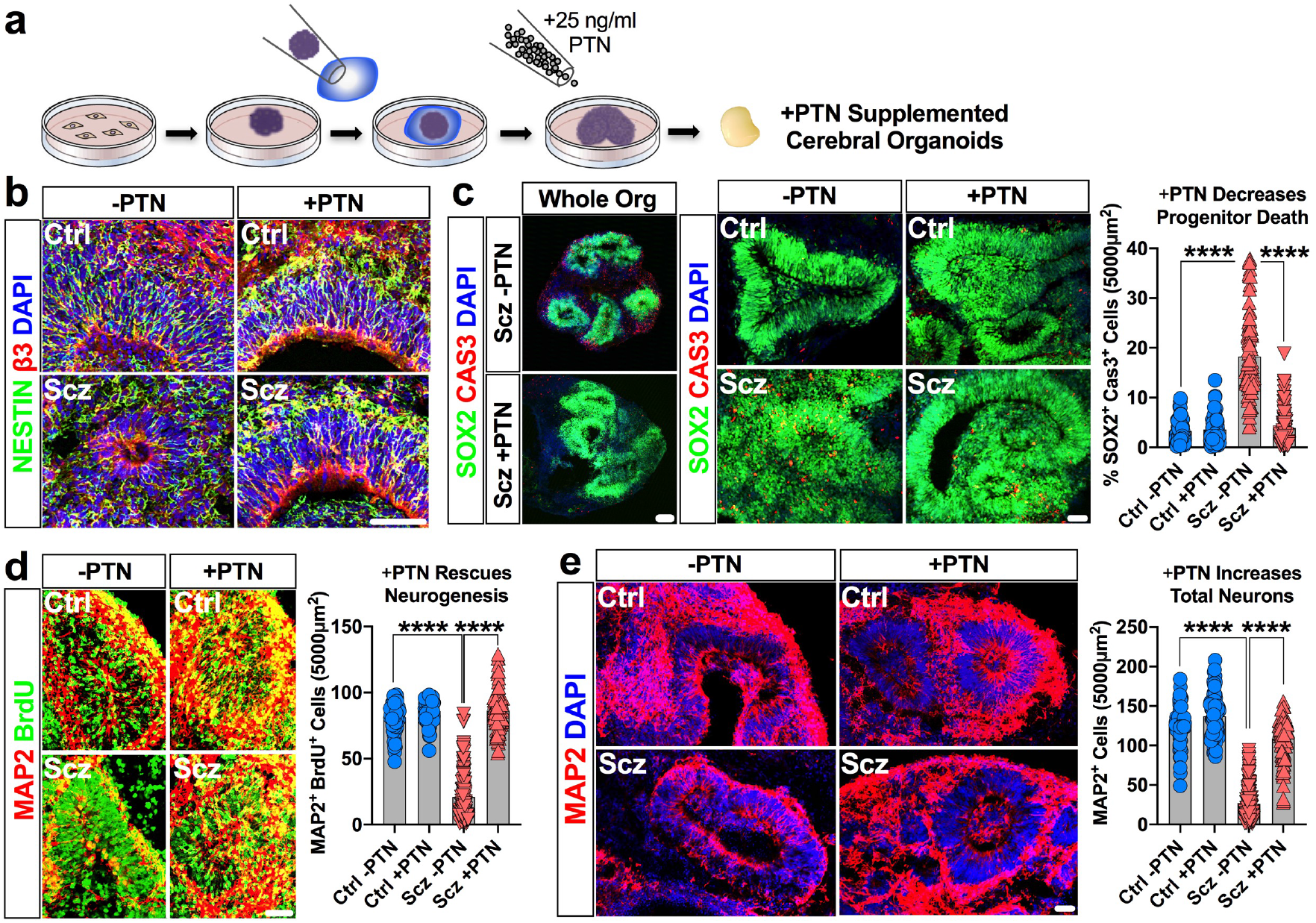
PTN rescues both survival and neurogenesis in Scz Organoids. a, Schematic of PTN-supplementation regime for rescue experiments. Proteomics revealed a decrease in total PTN expression in Scz organoids (fig. 2e). Similarly, scRNA-Seq revealed that Scz organoids exhibited lower PTN gene-expression in progenitors and neurons (fig. 3m-n). We identified that organoids supplemented with 25ng/ml PTN (a concentration for growth factor supplements) resulted in viable tissue without gross morphological evidence of deterioration. This subsequently permitted a laboratory-controlled examination of PTN’s role in intrinsically arising neuropathology of Scz in self-developing human-derived neural tissue. **b-c, PTN treatment promotes progenitor survival in Scz organoids.** PTN is expressed in most cell type clusters in organoids (fig. 3m) but is particularly enriched in neural progenitors (fig. 3n). Post treatment, the ventricular zones of PTN- supplemented Scz organoids often appeared enriched for Nestin (b). PTN treatment rescued progenitor death in the ventricular zones of Scz organoids to levels consistent with Ctrl cultures (panel c; Vehicle Ctrl *n* = 69 fields, *n* = 28 organoids, and *n* = 5 independent Ctrl lines; PTN-supplemented Ctrl *n* = 70 fields, *n* = 30 organoids, and *n* = 5 independent Ctrl lines; Vehicle Scz *n* = 90 fields, *n* = 44 organoids, and *n* = 9 independent Scz lines; PTN-supplemented Scz *n* = 85 fields, *n* = 41 organoids, and *n* = 9 independent Scz lines). Enlarged, whole-organoid, images are also provided in the Supplementary Material (see fig. S9). PTN treatment also broadly increased new-born cell survival in Scz organoids (see fig. S10). These data suggest that PTN exerts neurotrophic-like effects on neural progenitors in organoids, and that low PTN expression in Scz organoids regulates neural progenitor cell death. As with our prior mechanistic rescue figure, raw data are graphed here for phenotype transparency and each data point on graphs reflects an independent ventricular zone (see fig. S11 for averaged data). **d-e, PTN supplementation restores neuronal differentiation in Scz organoids.** We next sought to determine if PTN levels contribute to neurogenesis phenotype in Scz organoids. To do this, we time-locked our 24hr-7d BrdU pulse-chase paradigm to coincide with commencement of PTN treatment. As in previous experiments, Scz organoids exhibited disrupted neuronal differentiation relative to Ctrl organoids. However, PTN treatment rescued the disrupted neurogenesis of Scz organoids. PTN supplementation restored the number of MAP2+ BrdU+ neurons in Scz organoids to levels consistent with Ctrl cultures (panel d; Vehicle Ctrl *n* = 75 fields, *n* = 26 organoids, and *n* = 5 independent Ctrl lines; PTN-supplemented Ctrl *n* = 79 fields, *n* = 27 organoids, and *n* = 5 independent Ctrl lines; Vehicle Scz *n* = 130 fields, *n* = 45 organoids, and *n* = 10 independent Scz lines; PTN-supplemented Scz *n* = 106 fields, *n* = 42 organoids, and *n* = 10 independent Scz lines). Thus, in addition to promoting progenitor survival, PTN- supplementation also promotes neuronal differentiation in Scz organoids. Lastly, and consistent with all prior phenotypes, PTN treatment increased total MAP2+ neuron numbers in Scz cortical fields as expected (panel e; Vehicle Ctrl *n* = 58 fields, *n* = 19 organoids, and *n* = 4 independent Ctrl lines; PTN-supplemented Ctrl *n* = 67 fields, *n* = 25 organoids, and *n* = 4 independent Ctrl lines; Vehicle Scz *n* = 150 fields, *n* = 57 organoids, and *n* = 11 independent Scz lines; PTN-supplemented Scz *n* = 97 fields, *n* = 41 organoids, and *n* = 10 independent Scz lines). Together, these experiments established that PTN mechanistically contributes to neuron numbers in 3D cortical assemblies within Scz organoids. Each data point on graphs reflects an independent, non-overlapping, cortical field, for phenotype transparency, with averaged data provided in Supplementary Material (see fig. S11). \*\*\*\**p* < 0.0001. Error bars reflect Standard Error of the Mean. Scale bar: 60µm. Ctrl: Control, Scz: Schizophrenia.

## DISCUSSION

The idea that Scz is defined by developmental pathology has been widely discussed in the literature, however the timeline for when Scz-related pathology commonly commences has remained speculative. This is largely because of an inability to both access and ethically study developing human neural tissue *in utero*. Here we resolve, for the first time, that Scz is defined by cell-specific neuropathology and that *multiple* mechanisms operate within self-developing patient-derived neural tissue of human origin. We report that a series of downstream alterations in Scz neuronal progenitors ultimately yielded a depletion of Scz neurons within Scz patient-derived organoids, which is indicative of neurogenesis defects similar to those reported in several animal models, 2D iPSC studies from schizophrenia donors, and postmortem analyses.

In our organoid cultures, we first identified that the ventricular progenitor pool was defined by increased rates of cell death. In further experimentation (fig. 1e and 3), we identified that Scz progenitors exhibited disrupted differentiation and did not produce neurons in expected quantities (fig. 1e and 2h-i). In single-cell analyses, we determined that Scz organoids exhibited cell-specific reprogramming in both progenitors and neurons (fig. 3d-l), that ultimately resulted in misdirected progenitor differentiation (fig. 3a-g) and reduced growth factor support (fig. 3h-l). This single-cell analysis identifies, for the first time, that neuropathology of Scz is vastly more complex than believed as disease signatures are likely to be encoded in a cell-specific manner within cellular lineages (e.g. progenitors vs. neurons), leading to both similarities but also distinct differences between cell-types. Thus, different cellular identities may encode and exhibit unique and non-overlapping disease pathology of Scz (fig. 3), leading to neuropathology that is encoded on a cell-by-cell basis.

Importantly, because cerebral organoids recapitulate *in utero* brain development, these experiments also establish that cell-specific mechanisms of disease are also likely to converge upon common mechanisms and/or pathways important for early brain development. Thus, while different cell-types may exhibit unique neuropathology of disease, our data also indicate that some similarities between cell-types are also likely to exist. Notably, we identified that both Scz progenitors and Scz neurons exhibited discrete growth factor alterations within the neurotrophin system family (fig. 3l).

Additionally, we identified a gross depletion of BRN2 and PTN in Scz within progenitors and neurons (fig. 3m-n). Importantly, in two separate rescue experiments, we exemplified that neuron numbers could be modulated when these mechanistic substrates are reconstituted back into Scz organoids. However, while BRN2 could modulate neurogenesis within Scz organoids, it could not rescue other Scz phenotypes (e.g. progenitor apoptosis; fig. 4). Notably, PTN was putatively able to restrain apoptosis while simultaneously promoting neuronal development (fig. 5). When amalgamated, these data also identify that ensembles of endogenously disrupted disease factors are capable of eliciting multiple potential mechanisms during early brain development.

Mechanisms of Scz have traditionally been studied within the adult nervous system. This is due to the protracted age of onset of Scz and relatively low incidence. However, there is also increasing debate whether disease phenotypes represent a ‘*cause or consequence’* of disease in the adult brain. Meanwhile, for several decades, evidence has continued to accumulate that early brain development is a critical period for Scz risk. Yet, without a way to ethically acquire, study, and subsequently identify whether human fetal tissue would definitively become a future Scz case, little progress has been made in understanding early developmental mechanisms of Scz. In this respect, patient-derived cerebral organoids are technically and ethically a model system for studying self-developing *human* tissue in 3D. Notably, cerebral organoids recapitulate the earliest phases of brain development, including corticogenesis. Indeed, our scRNA-Seq revealed that Ctrl organoids predominantly comprised progenitors, proliferating cells, and developing cortical cell-types as expected (fig. 2). TMT-LC/MS supported this by exemplifying widespread induction of neural ontology pathways required for brain development (see Supplementary Tables). Across a number of experiments, we identified critical differences between Ctrl and Scz organoids within the ventricular progenitor pool. Relative to Ctrls, we identified that ventricular progenitors in Scz organoids exhibited increased apoptosis and decreased neuronal differentiation. Critically, in a cell-fate pulse-chase experiment (fig. 1e), we identified that Scz progenitors exhibited a 3-fold decrease in neurogenesis and a 2-fold increase in rates of differentiation into “other” neural-related cell-types. In a follow-up analysis of 26,335 single-cell transcriptomes, we identified the identity of these other cell-types. Namely, we found that an enrichment for neuroendothelial and neurovascular-related cell-types that, despite of their differences in lineage and ontogenesis, were seemingly produced at the expense of neurons (fig. 2-3). In deeper analyses, we identified that Scz progenitors exhibited a reprogramming in their expression of neuronal differentiation markers (fig.

3d-g), including transcription factors such as BRN2 that are almost completely diminished (fig. 3m-n). Other altered factors in Scz progenitors comprised neuronal development factors, such as HOXA2, AUTS2, SLITRK6, WNT2B, NRN1 and NT-3, which also collectively orchestrate neuronal differentiation (fig. 3e). Instead of exhibiting these expected cell-fate programming factors, Scz progenitors exhibited enrichment for extracellular matrix factors, actin filaments, cell-substrate adhesion, vascular-related metabolic processes, and angiogenic markers (fig. 3). In pseudotime trajectories, these factors predicted the altered differentiation trajectories of Scz progenitors away from neurogenesis and towards different cellular lineages (specifically, mural, myeloid, and endothelial cells; see fig. 3a-b). While the numbers of these cell-types were abnormal in Scz organoids, it is nonetheless important to emphasize that these cells are necessary for brain development and are ordinarily found in Ctrl organoids too (fig. 3h). Therefore, in sum, computational modeling consequently identified the induction of altered cell-lineages in Scz organoids and the depletion of an ensemble of neuronal programming factors in Scz progenitors. This outlines a suite of novel neuropathology in developing human tissue that sheds light on early-arising alterations in the developing cerebral cortex that may be related to intrinsic Scz risk.

As discussed above, a critical finding in our manuscript was the observation that Scz progenitors seemingly lacked the transcriptional machinery necessary to support neuron production and thus neocortical neurogenesis. To provide mechanistic support for this hypothesis, we sought to identify a neurogenic target that could be exogenously modulated *in situ* within organoids. This led us to focus on *BRN2* (*POU3F2*), as Scz organoids exhibited robust down-regulation of this factor at the mRNA level as well as POU-domain fragments at the peptide level (fig. 2e and 3m-n). Prior studies have implicated that BRN2 is important for the induction of progenitor differentiation into neurons within the developing cortex [71]. Additionally, in a prior study it was reported that disruption of DISC1 resulted in elevated progenitor mitotic activity and altered BRN2+ neuron numbers [72], which mimicked the phenotypes reported here. This led us to target BRN2 in our first rescue experiment, whereby we utilized a self-regulating viral vector to deliver exogenous BRN2 selectively to Scz progenitor cells but not neurons.

This led us to confirm that supplementation of BRN2 levels in Scz progenitors reinstated their neurogenic potential, yielding increased neuron numbers in Scz organoids (fig. 4). Thus, with no major disease association at the genetic level (fig. S6), these experiments established BRN2 as a forebrain-specific transcription factor that may contribute to Scz neuropathology as a downstream intermediary during early brain assembly. However, because BRN2 did not rescue Scz progenitor apoptosis, it was also apparent that yet further mechanisms were operating within our 3D Scz patient-derived organoids.

In addition to neuronal programming factors, Scz progenitors were further defined by cell-specific alterations in the expression of growth factors. This was of note, as growth factors regulate not only neurogenesis but also cellular survival. Growth factor alterations included the significant down-regulation of *NT-3* (*NTF3*) and *TrkB* (*NTRK2*) gene-expression in Scz progenitors (fig. 3l). Gene variants within NT-3 [97], and epistatic interactions between *BDNF* and *TrkB* gene variants [98], have previously been suggested to precipitate Scz risk. Intriguingly, functions linked to these neurotrophins were also hallmark neuropathology of Scz organoids. Indeed, neurotrophins and TrkB-mediated signaling are potent regulators of differentiation and survival [99–101] and likely elicit neuroprotective effects in Scz [102]. Neurotrophins also orchestrate transcriptional mechanisms of neuronal fate making [103–106] and protect neural stem cells from apoptosis via activation of Pl3K/MAPK signaling [100]. A disruption in growth factor homeostasis within Scz progenitors may therefore contribute to alterations in disrupted neurogenesis and increased progenitor cell death. Interestingly, *NT-4* expression was selectively up-regulated in Scz progenitors, and Scz neurons exhibited none of these alterations but instead exhibited a ∼3-fold depletion in *TrkA* mRNA expression. These cell-specific differences are thus consistent with our suggestion of discrete cell-specific neuropathology of cell-types belonging to the same overall cellular lineage that converge upon common pathways and/or targets. Thus, the convergence of both Scz progenitors and neurons upon neurotrophin dysregulation support this hypothesis. Additionally, the divergence in expression between these cell-types likely reflect differences in the temporal coordination and cell-specific actions [107] of individual neurotrophins [108, 109] during brain development. These data correspondingly evidence the cell-type specificity of growth factor neuropathology in Scz, and identify that a developmental switch in the expression of neurotrophin-related factors occurs as progenitors differentiate into neurons.

Disrupted neurotrophin homeostasis within the progenitor pool led us to target a putative growth factor for our second rescue experiment. Similar to our BRN2 rescue experiment, we assigned priority to factors that exhibited independent cross-validation at both the mRNA and peptide level within Scz organoids. This selection criterion led us to target PTN for our second rescue experiment. PTN was depleted in Scz progenitors and neurons (fig. 3m-n) as well as in our proteomics assays (fig. 2), and PTN also functions as a potential embryonic growth factor [110] in the developing cerebral cortex [111].

Evidence supporting a genetic association between PTN and Scz is defined by an index SNP in proximity to PTN that was identified in the largest-ever Scz GWAS (*n* = ∼150,000) [36] (see also fig. S6). However, because this comprised the discovery of a novel locus, the role of PTN in Scz has remained unclear and thus undefined. Here we discovered that PTN exerted neuroprotective effects in Scz organoids by restraining apoptosis whilst simultaneously promoting neurogenesis (fig. 5). Therefore, we also mechanistically established that PTN positively regulated neuron numbers in Scz patient-derived organoids albeit via similar, yet independent, effects to BRN2 (see discussion below). Overall, this experimentally established that multiple Scz mechanisms are likely to be operational during corticogenesis within patient-derived organoids.

It is important to emphasize that our rescue experiments established that BRN2 and PTN rescue Scz neuron numbers via related, but distinctly different, effects. This principally included neuronal differentiation and survival. In fig. 4-5, we exemplify that both neuronal differentiation and survival factors are likely to be mutually important, and that disease-implicated factors likely amalgamate to alter neurodevelopment within Scz organoids. This is broadly supported by our Scz phenotypes described throughout the manuscript, including a depletion of normative neuronal development factors in progenitors (fig. 3e-g) that potentially divert progenitor cellular lineages away from neurogenesis (fig. 1e and 3a-c). Additional correlates included endogenously diminished growth factor support (e.g. fig. 2e, 3l, 3m-n, 5), increased apoptosis (fig. 1, S2, 4, and 5), and altered differentiation trajectories (fig. 1e and 3a-c) in Scz organoids. Indeed, we were able to repetitively replicate progenitor death and neuronal depletion (fig. 1, 2, 4, and 5). However, as shown in our two independent rescue experiments (fig. 4-5), disrupted neuron production could only be reinstated if mechanistic substrates (e.g. BRN2 or PTN) were reconstituted in Scz organoids. Because BRN2 functions as a differentiation-inducing transcription factor and PTN as a growth factor, this suggests that alterations in neurogenesis and survival factors yielded an intrinsically ligated potential for cortical development within Scz organoids (figure 1e, 2h-i, 3a, 4, 5). Consistent with this, complete phenotypic rescue could only be achieved if progenitor apoptosis was normalized alongside neurogenesis; as was the case only in our PTN rescue experiments (fig. 5) but not our BRN2 rescue experiments (fig. 4). This disparity between our mechanistic rescue experiments exemplifies that numerous molecular agents are capable of regulating similar, yet also distinct, Scz neuropathology within patient-derived organoids, and that neurogenesis defects alone do not explain the breadth of Scz organoid phenotypes. Instead, these data support that mechanisms regulating cell death and neuronal differentiation are dysfunctional in self-developing Scz patient-derived organoids, resulting in a terminal upstream depletion of cortical neurons. This supports the idea that multiple mutual mechanisms operate within Scz organoids, and that Scz disease factors may target similar biological processes and pathways necessary for neurodevelopment.

While our work provides proof of principle that multiple factors regulate intrinsic neuropathology of Scz organoids, we believe that yet more mechanisms are likely to be operational in Scz organoids. Importantly, alterations in the availability of other Scz molecular factors may also produce differential or synergistic risk [112]. It will be particularly important to parse the functionality of Scz risk factors validated in both proteomics and scRNA-Seq. A prominent target may be another novel Scz risk factor with nascent genome-wide significance [36], namely PODXL (a neural adhesion molecule involved in axonal fasciculation, neurite outgrowth, and synaptogenesis [76]). Other targets included longstanding Scz risk factors such as PLCL1 (involved in GABAergic neurotransmission [74]) and COMT (involved in dopamine and catecholamine elimination [77] and cognitive response to antipsychotics [78]). These factors are therefore potential targets for unraveling yet further mechanisms of Scz risk.

Our unbiased datasets also identified several other potential disease factors that do not hold genetic association with Scz but exhibit face validity for future investigations. Namely, our data also revealed an intrinsic enrichment for the interferon-induced transmembrane factor *IFITM3* in Scz organoids. *IFITM3* enrichment was present across a multitude of cell-type clusters. In the postmortem Scz brain, enrichment of *IFITM3* in the hippocampus [113], amygdala [114], and frontal cortex [115] has been reported and may reflect increased neuroinflammation. These phenotypes may be related to increased markers of oxidative stress in Scz neural progenitors [95]. Interestingly, IFITM3 has also been suggested to mediate perinatal/neonatal immune activation effects on the brain [116] and, consistent with this, knockout mice are protected from neuronal pathology induced by perinatal Poly(I:C) treatment [82]. IFITM3 has also been implicated in another disorder defined by immune-related risk during neurodevelopment, namely autism [117]. Correspondingly, IFITM3 has even been proposed as a novel drug target for Scz [118]. These data only enhance the novelty of our discovery that *IFITM3* is enriched in Scz patient-derived organoids and suggests that neuroinflammation may correspondingly be an intrinsic Scz phenotype during early brain development. In support of this, our data also identified cell-specific up-regulation of the interleukin-6 signal transducer *IL6ST* (or, *GP130*) specifically within Scz neurons. Given that IL6ST mediates the induction of downstream transcriptional effectors of IL-6, and was associated with growth factor binding factors in Scz neurons (fig. 3i-j). This result was therefore also intriguing. Similar to *IFITM3,* there is preliminary evidence that *IL6ST* holds putative disease relevance to Scz. This includes elevated *IL6ST* expression in Scz postmortem midbrain samples [119] as well as altered expression in the dorsolateral prefrontal cortex of a mixed postmortem cohort with high neuroinflammation [121]. Interestingly, *IL6ST* in the prefrontal cortex may only be associated with shorter durations of illness [120]. This implicates that *IL6ST* may be involved in early disease induction and/or states, which is consistent with its emergence in our neurodevelopmental dataset. Cumulatively, these data provide fundamental support for the idea that inflammation may be an intrinsic component of early developmental neuropathology in Scz patient-derived organoids. Therefore, *IFITM3* and *IL6ST* represent fruitful targets that are deserving of further study within the context of Scz and relevant neurodevelopmental hypotheses.

Lastly, an important avenue of further research will be to further map the functional qualities of various cell-types identified in cerebral organoids. For example, single-cell analysis of neuronal dynamics via calcium signaling or more elaborate single-unit electrophysiology may provide important functional data on Scz neuron activity within a 3D macroenvironment. It is also plausible that other organoid models, other cell-types, or specific patient groups may yet unveil disease effects that were not identified here. In this respect, just as we hypothesize that more mechanisms of Scz exist in Scz organoids, we predict that more Scz phenotypes also remain to be discovered. It is therefore plausible that greater sampling may be required to identify subtle phenotypes, or that clinically stratified samples are required to identify how certain Scz cases segregate between clinical or genetic Scz subtypes. While these experiments are outside the scope of the current study, future studies should consider the utility of other single-cell technologies and develop pipelines that enable a broader range of patient samples to be functionally interrogated.

## CONCLUDING REMARKS

In closing, our patient-derived organoid data conceptually validates the principle behind numerous long-held hypotheses regarding the developmental basis of Scz. We experimentally establish that an ensemble of novel Scz risk factors likely amalgamate in various combinations to trigger the ontogenesis of Scz developmental neuropathology. We specifically exemplify via unbiased deep-phenotyping that organoids provide utility by enabling Scz mechanisms to be parsed in a *human* 3D environment that also captures the broader molecular-genetic background of disease. Our study also identifies the disease relevance of two novel factors, PTN and BRN2, in Scz organoids.

Importantly, we establish that these factors positively regulate neuronal development in Scz patient-derived organoids via similar but putatively distinct mechanisms. While reconstitution of BRN2 was shown to only regulate neurogenesis deficits in Scz organoids (fig. 4), PTN modulated death differentiation, and neuron numbers within Scz patient-derived organoids (fig. 5). This indicates that multiple mechanistic factors can contribute to both independent phenotypes and convergent (i.e. common) disease pathology.

This work therefore defines novel early-arising signatures of Scz at the single-cell and posttranslational levels in 3D organoids. When amalgamated, these data therefore suggest that multiple cell-specific mechanisms of Scz likely exist, which converge upon primordial brain developmental pathways such as neuronal differentiation, survival, and growth factor support, to elevate intrinsic risk of Scz within the early developing brain.

## MATERIALS AND METHODS

### Human induced pluripotent stem cells

All human iPSC lines were sourced from NIH deposits at the Rutgers University Cell and DNA Repository. All lines have undergone extensive characterization for identity, pluripotency, exogenous reprogramming factor expression, genetic stability and viability. A total of 11 different Induced Pluripotent Stem Cell (iPSC) lines, each coming from a unique human donor, were utilized between organoid experiments. A comprehensive list and description of iPSC donors and their clinical characteristics are provided in Supplementary Table 1. All Scz samples were derived from idiopathic cases, which we define here as schizophrenia cases that maintained unknown disease origins and do not meet a genetic/syndrome-based diagnosis that explains disease. A breakdown of idiosyncratic iPSC properties, in the context of organoid generation, maturation, survival and subsequent experiment inclusion details, can be found in Supplementary Table 2.

All iPSC lines were maintained on vitronectin-coated plates and fed with Essential 8 (E8) + E8 supplement media (ThermoFisher, CAT#: A1517001).

### Three-dimensional cerebral organoid culturing system

We adapted the undirected-differentiation organoid system published by Lancaster et al. in *Nature* [1] and *Nature Protocols* [2]. While some discussion of this system was included in the manuscript, allowing the context of results to be framed by a broader audience, this culturing system can be minimized into four major stages. First, iPSCs are dissociated with Accutase (Laboratory Disposable Products, CAT#: 25-058-C1) and cultured into three-dimensional embryoid bodies for up to 7 days using previously described media (see [2]) in ultra-low attachment 96 well plates (Corning; CAT#: 3474). Rock inhibitor (1:1000; Stem Cell Tech, CAT#: 72304) and basic fibroblast growth factor (Pepro Tech, CAT#: 100-18B) are included in media for the first 2-4 days of embryoid body culturing to promote stem cell aggregation and survival. Following this, healthy embryoid bodies are isolated and transferred to Nunclon Sphera 24 well plates (Thermo Scientifiic, CAT#: 174930) for neural fate specification, using custom neural induction media (see [2]). Once neuroepithelium was apparent (see Supplementary Figure 1), successful early ‘organoids’ were embedded in a 30µl Matrigel (Corning, CAT#: 354234) spheroid-droplet and polymerized at 37°C for 20-30min which provided a matrix for subsequent neural expansion. Organoids suspended in matrigel droplets were next cultured in terminal organoid media (see [2]) for 4-6 days without agitation, and then cultured with agitation at 60-70RPM until harvested for experiments. All stages of culturing occurred at 37°C with 5% atmospheric CO2 in a sterile incubator. Cerebral organoids from healthy control and schizophrenia iPSC lines were generated and maintained in parallel, ensuring that idiosyncratic differences in culturing conditions were accounted for between control and schizophrenia organoids. Lastly, all cultures underwent quality control assessments on a rolling basis, utilizing the published criterion and guidelines in [2] which describes numerous “Go/No Go” criteria at different stages of organoid generation. Additionally, inclusion/exclusion criterion were adapted from other organoid protocols which comprised screening for evidence of neural induction including the presence of neural stem cells/progenitors (SOX2+, NESTIN+ cells), forebrain/cortical-specific progenitors (FOXG1+, PAX6+ cells), and early-developing neurons (DCX+, MAP2+ cells) including those with cortical-specific identities (CTIP2+, SATB2+, and BRN2+ cells). This ensured that all lab-generated tissue in the manuscript exhibited neural induction as well as cell-types specific for the developing cortex. Independent confirmation of neuronal induction was also achieved via unbiased analysis of our scRNA-seq and proteomics datasets.

### Immunohistochemistry and laser-scanning confocal microscopy

Prior to immunohistochemistry, organoids were drop-fixed in 4% paraformaldehyde, dehydrated in 30% sucrose, embedded in Tissue-Tek OCT compound (CAT#: 4583) using biopsy molds and cryosectioned serially *between-slides* at 30µm (∼25 slides per organoid). Each slide therefore contained 3-4 unique sections/Fields of View (FOV) from each organoid studied. This allowed for robust assessment of both independent and focal cell populations in each biological and technical organoid replicate. All sections underwent heat-mediated antigen retrieval in citrate buffer, and incubated in primary overnight. Primary antibodies comprised SOX2 (1:1000; R&D Systems, CAT#: MAB2018-SP), PAX6 (1:1000; Biolegend, CAT#: 901301), β-tubulin III (1:1000; Abcam, CAT#: AB41489), MAP2 (1:1000, Abcam, CAT#: AB11267; 1:1000, Abcam, CAT#: AB32454), BrdU (1:1000, BD Pharminogen, CAT#: 555627), DCX (1:1000, Santa Cruz, sc-271390), Activated/Cleaved Caspase-3 (1:1000; Cell Signalling Biology, CAT#: 9661S and Thermo Fisher Scientific, CAT#: 66470-2-IG), Nestin (1:1000, Abcam, CAT#: AB176571), pH3 (1:1000; Millipore, CAT#: 06-570), Ki67 (1:1000; BD Biosciences, CAT#: 550609), TBR2 (1:300; Abcam, CAT#: AB23345 and EMD Millipore, CAT#: AB15894), GFP (1:1000, Thermofisher, CAT#: A10262), CTIP2 (1:300; Abcam, CAT#:

AB18465), and BRN2 (1:750, Santa Cruz, CAT#: SC-393324). Secondary antibodies were incubated for 2 hours at room temperature, and comprised antibodies for rabbit (Fluor 488 CAT#: A11008; Fluor 546 CAT#: A11035; & Fluor 633 CAT#: A21070), mouse (Fluor 488 CAT#: A11001; Fluor 546 CAT#: A11003; & Fluor 633 CAT#: A21052) and chicken (Fluor 546 CAT#: A11040) were used at a 1:2000 dilution, and sourced from Life Technologies. Microscopy was completed on an Olympus IX81 Laser-Scanning Confocal Microscope, controlled by proprietary Olympus FluoView software. Images were typically acquired at 1200x1200 resolution with optical *Z* slices (step sizes) ranging from 0.5-10µm depending on the unit of analysis.

### BrdU pulse-chase assay for mapping cell fate

Neuronal differentiation experiments comprised a 24hr pulse of BrdU and a 7d chase period. Organoids were pulsed with 10µM BrdU in media that was aspirated 24hr later. Organoids were washed three-times with differentiation media and left to maturate.

Approximately 7d following BrdU pulse, organoids were drop-fixed, cryosectioned, and immunostained to examine progenitor self-renewal and neuronal differentiation (fig. 1).

### High-throughput flow cytometry of apoptosis & DNA damage in organoids

To derive an unbiased assessment of DNA damage, cell death, and DNA-damaged cells undergoing apoptosis following drug treatment, we adapted a well validated and widely utilized FACS kit (BD Pharmingen, CAT#: 562253). Briefly, pseudorandomly selected organoids were dissociated to a single-cell suspension via a 20min exposure to accutase followed by tricheration and serial filtering through 70→30μm pores. Cells were resuspended and incubated in in Cytofix/Cytoperm solution for fixation and initial permeabilization for 30min at room temperature. Cells were consequently washed, resuspended in BD “Plus” permeabilization buffer for 10min on ice, and re-exposed to Cytofix/Cytoperm solution for 5 additional min to achieve refixation. Cells were consequently washed, and resuspended in 30μg of DNAse for 60min at 37°C. Cells were consequently washed and labeled with PE-Cleaved PARP (BD Pharmingen, CAT#: 51-9007684) to label cells committed to apoptosis and Alexa647-H2AX (pS139, BD Pharmingen, CAT#: 51-9007683) for cells exhibiting DNA damage. Double positive cells represented apoptotic cells exhibiting DNA damage. Per manufacturer instructions, antibodies were diluted at a 1:25 ratio and incubated for 20-minutes at room temperature. Cells were washed and resuspended in staining solution. Labeled suspensions were analyzed utilizing a BD Aria II (Becton Dickinson) cell sorter to acquire multiparameter data files. Data were presented as a % of H2AX+ DNA damaged cells, and % of PARP+ H2AX+ double-positive cells that exhibited both DNA damage and induction of cell death.

### Single-cell DNA Content Analysis

Organoids were prepared to a single-cell suspension as described above, and washed successively with calcium/magnesium free PBS at 4°C to remove residual peptides in solution. Samples were consequently centrifuged, supernatant removed, and pelleted cells resuspended in PBS. Cells were EtOH fixed (100%, at 4°C) while being gently vortexed. Cells were rehydrated, and incubated with Triton-X with DNAse added for 5 mins. Cells were pelleted, Triton-X removed, and resuspended in 200μl of calcium/magnesium free PBS at 4°C. Immediately prior to flow cytometry, suspensions were incubated with 2μl of Propidium Iodide (PI; Thermofisher, Material#: P3566), and analyzed for single-cell DNA content utilizing a BD Aria II (Becton Dickinson) cytometer. Analysis was subsequently modeled *post hoc* in the FloJo cytometry analysis package (Becton Dickinson).

### Tandem Mass Tag (TMT) Liquid Chromatography-Mass Spectrometry (LC/MS)

TMT mass-spectrometry proteomics was completed as previously described [6]. Briefly, organoids were reduced with dithiotreitol and underwent alkylation with iodoacetamide before tryptic digestion at 37°C overnight. Peptide suspensions were desalted using C18 stage-tips prior to Liquid Chromatography-Mass Spectometry (LC-MS) analysis. An EASY-nLC 1200, which was coupled to a Fusion Lumos mass spectrometer, (ThermoFisher Scientific) was utilized. Buffer A (0.1% FA in water) and buffer B (0.1% FA in 80% ACN) were used as mobile phases for gradient separation [6]. A 75µm I.D. column (ReproSil-Pur C18-AQ, 3µm, Dr. Maisch GmbH, German) was packed in-house for separating peptides. A separation gradient of 5–10% buffer B over 1min, 10%-35% buffer B over 229min, and 35%-100% B over 5min at a flow rate of 300nL/min was adapted. Data dependent mode was selected during operation of the Fusion Lumos mass spectrometer. An Orbitrap mass analyzer acquired Full MS scans over a range of 350-1500m/z with resolution 120,000 at m/z 200. The top 20 most-abundant precursors were selected with an isolation window of 0.7 Thomsons and fragmented by higher-energy collisional dissociation with normalized collision energy of 40. The Orbitrap mass analyzer was also used to acquire MS/MS scans. The automatic gain control target value was 1e6 for full scans and 5e4 for MS/MS scans respectively, and the maximum ion injection time was 54ms for both.

### Bioinformatics pipeline for TMT-LC/MS proteomics

Mass spectra were pre-processed as described [7] and processed using MaxQuant [8] (1.5.5.1). Spectra were searched against the full set of human protein sequences annotated in UniProt (sequence database Sep-2017) using Andromeda. Data was searched as described [9] with fixed modification, cysteine carbamidomethylation and variable modifications, N-acetylation and methionine oxidation. Peptides shorter than seven amino acids were not considered for further analysis because of lack of uniqueness, and a 1% False-Discovery Rate (FDR) was used to filter poor identifications at both the peptide and protein level. Protein identification required at least one unique or razor peptide per protein group. Contaminants, and reverse identification were excluded from further data analysis. Resulting *p* values were adjusted by the Benjamini-Hochberg multi-test adjustment method for a high number of comparisons [10] and statistics performed as previously described [11]. The proteome profiles from the analyses of two biological replicates and individual technical replicates were combined by calculating the average intensity. Proteins that were quantified in only one replicate were also included in the combined data set. For further data analysis, the normalized intensities were converted into log2 ratios of the intensities over the median intensity measured for each protein across each sample group.

For enrichment and pathway analyses, Kyoto Encyclopedia of Genes and Genomes (KEGG) and NIH Database for Annotation, Visualization and Integrated Discovery Bioinformatics Resources 6.7 (DAVID) platforms were utilised using recommended analytical parameters [12]. For gene ontology (GO) enrichment and network analyses UniProt (www.uniprot.org, [13]) database resource (biological process, molecular function), Ingenuity Pathway Analysis, and Reactome knowledgebase were utilized. Clustering of sample groups was performed by principal component analysis (PCA) and visualized using ggplot2 [14] and ggfortify (https://cran.r-project.org/web/packages/ggfortify/index.html; [15]). Protein abundance distribution for global and subset groups was performed using gplots (https://cran.r-project.org/web/packages/gplots/index.html; [16]. Based on hypergeometric distribution, the GO terms and pathways that differentially expressed proteins are significantly enriched-in were calculated. Benjamini-Hochberg method was used to correct obtained *p* values. A corrected *p* value ≤ 0.05 was considered to represent significant enrichment. Protein–protein associations were evaluated for differentially expressed protein subsets using STRING database. Correlation pairs were ordered, and redundancies were removed before comparison to a non-redundant version of the STRING database to find STRING-annotated interactions. Only STRING interactions of high confidence (score ≥ 0.700) were considered for proteome-target analyses.

Unbiased computational analysis of proteomics data utilized correction methods such as FDR rates and *p* value adjustment using the Benjamini-Hochberg method were adapted, as described above, and Statistical testing of was performed using a Poisson distribution with EdgeR software v3.2.

### 10x Chromium^TM^ microfluidic device single-cell RNA-sequencing

Single-cell sequencing was conducted as described in-text, figure legends, and depicted in our schematic pipeline (see fig. 2f). Briefly Organoids were dissociated using Accutase, serially filtered to remove debris, and were subjected to high-throughput FACS (Aria II flow cytometer, Becton Dickson). This procedure allowed us to isolate only live cells in a defined concentration (2000 live cells/µL) that could be standardized between lines for microfluidic-device loading. Cell viability was confirmed using Countess-II (Invitrogen, Thermofisher, CAT#: AMQAX1000). Next, cells were prepared for 10X Genomics Chromium library preparation according to manufacturer instructions (see Chromium^TM^ Single-Cell 3’ Library and Gel Beads and Chromium^TM^ CHIP kit, 10x Genomics). Post sample generation, barcoded-emulsions were broken, amplified, and libraries prepared. cDNA was examined on a Fragment Analyzer running PROSize v3.0 3.0.1.6 (Advanced Analytical Technologies). Libraries were subsequently subjected to next generation sequencing via Next-Seq High-Output 150-cycled 26-8-98.

### Bioinformatics pipeline for single-cell RNA sequencing

Initial processing of raw reads involved alignment to GRCh38 with the CellRanger v3.0.2 pipeline (10x Genomics, USA). Subsequent analyses were performed in R and Bioconductor (https://osca.bioconductor.org/) per guidelines [17]. This comprised numerous functions in the scater and scran packages [18]. Cells with fewer than 317 genes and more than 8% mitochondrial reads were removed based on outlier calculations. Subsequent processing and integration of samples was completed using Seurat v3.1 following vignette recommendations. More specifically, read counts were first individually normalized using SCTransform for each sample [19]. The different samples were then integrated using the top 3,000 most informative genes [20] before performing various dimensionality reduction steps including principal components analysis (PCA) and UMAPs [21]. Batch correction was also applied. A shared nearest neighbor graph was constructed with default settings (e.g. k = 20). This comprised the first 25 principal components. Subsequent clustering was performed with the resolution parameter set to 0.2. For visualizations and assessments of normalized expression values, SCTransform-normalized (log-transformed) expression values were principally adapted. Cell-type clusters were labelled based on marker enrichment, derived from both Seurat-identified clusters (using default parameters) as well as automated assignment with SingleR using reference datasets [22]. Seurat analysis comprised FindMarkers and FindAllMarkers functions were used with default settings as well as scran::FindMarkers(), comparing only cells of the control condition. For SingleR, we used its in-built reference data set of the human primary cell atlas, which contains RNA-seq from adult human tissues, including neurons and myeloid cells. In addition, we downloaded the single-cell RNA-seq data from numerous human fetal brain samples [23] and used it as the training data with the refactored bioconductor version of SingleR (http://www.bioconductor.org/packages/devel/bioc/html/SingleR.html).

After integration, we determined differential gene-expression using a pseudo-bulk approach [24] that summed the reads across all cells belonging to the same type of cells (where “type” would be determined by the sample and the membership within a given subpopulation of interest). To identify genes with significantly different expression (DEG) levels between two types of cells, we used methods implemented in the edgeR package including dispersion estimation and gene-wise negative binomial generalized linear models with quasi-likelihood tests [25]. Unless stated otherwise, DEG were based on adjusted p-values lower than 0.01. To test for over-representation of specific gene sets of KEGG, Reactome, and Gene Ontology collections in our DEG lists, we used the enrichment functions implemented in clusterProfiler and reactomePA [26] with an adjusted p-value cut-off of 0.05. Note, for STRING protein-protein interactome modeling of scRNA-Seq pathway factors (specifically, fig. 3k), specific analysis parameters were applied. Namely, kmeans clustering (*n* = 3) was applied, network edges were limited to molecular action, a minimum interaction confidence limit of 0.4 was adapted, and no more than 5 interactors in the first shell were allowed. Code used for scRNA-Seq analysis is available on github, and the dataset will be available via repository curation following publication of this manuscript.

### Diffusion maps and pseudotime trajectories of single-cell transcriptomes

For ordering the cells along pseudotime trajectories, we first constructed diffusion maps [27] that are implemented in the destiny package [28]. Cell lineages and temporally expressed genes were inferred using slingshot [29] following recommendations in vignettes and technical materials. In brief, pseudotime values were extracted for each cell along each lineage identified by slingshot. Then, each gene’s normalized expression values were regressed on the pseudotime variable using a general additive model to identify those whose expression patterns most closely correlate with the pseudotime weights. This allowed profiles of cellular differentiation to be computationally constructed in an unbiased environment based on large-scale single-cell sequencing data.

### Technical details of BRN2 and PTN rescue experiments

For BRN2 mechanistic rescue experiments, virus was administered during neural induction when 3D embryoid bodies remained at their lowest cellular densities. As described in-text, *CTRL*-Virus organoids were administered a GFP virus as a vehicle while *BRN2*-Virus organoids received a self-regulating BRN2-GFP viral vector (vector details, including complete viral schematic, are provided in Figure S10). As described in-text, this hPGK>BRN2:P2A:EGFP.MIRT124^4^ lentiviral vector was a GFP-modified version of a previously validated neuronal conversion tool [30]. Expression of the neuronal mRNA, miRNA-124, to four recognition units in virus transcripts (MIRT.124^4^) leads to translational suppression in neurons but not other cell-types [30]. Specifically, Expression of miRNA-124 represses *SOX9* to direct progenitors towards a post-mitotic neuronal fate [87] where miRNA-124 expression is preferentially maintained [88] and reaches its peak [89]. This high intracellular expression of miRNA-124 in neurons suppresses translation of *BRN2*-Virus, leading to complete loss of exogenous BRN2- GFP in mature post-mitotic neurons only. Organoids were infected under uniform conditions irrespective of line, preparation, or group allocation. Virus and polybyrene was supplemented into neural induction media for 24hr. Organoid generation then continued as described without further modification. For PTN mechanistic rescue experiments, organoids were also generated as described excepting the addition of 25ng/mL recombinant human PTN (Peprotech, CAT#: 450-15) into differentiation media for 7d.

This concentration was derived as it is similar to concentrations of morphogens used in other organoid studies, and this yielded viable 3D tissue without evidence of deterioration. Organoids allocated to the vehicle-treatment group received culture-grade diluent without PTN. For pulse-chase experiments in PTN-supplemented organoids, BrdU administration coincided with first exposure to PTN. Organoids were pseudorandomly selected to treatment groups for both mechanistic rescue experiments.

*Virus Sequence for POU3F2 [BRN2; NM_008899.2; 1335/9305bp)*

ATGGCGACCGCAGCGTCTAACCACTACAGCCTGCTCACCTCCAGCGCCTCCATCGTACATGCCGAGCCGCCTGGCGGCATGCAGCAGGGCGCAGGGGGCTACCGCGAGGCGCAGAGCCTGGTGCAGGGCGACTACGGCGCGCTGCAGAGCAACGGGCACCCGCTCAGCCACGCTCACCAGTGGATCACCGCGCTGTCCCACGGCGGCGGCGGCGGGGGCGGCGGCGGCGGTGGAGGAGGCGGGGGAGGCGGCGGGGGAGGCGGCGACGGCTCCCCGTGGTCCACCAGCCCCCTAGGCCAGCCGGACATCAAGCCCTCGGTGGTGGTACAGCAGGGTGGCCGAGGCGACGAGCTGCACGGGCCAGGAGCGCTGCAGCAACAGCATCAACAGCAACAGCAACAGCAGCAGCAGCAGCAGCAGCAGCAGCAGCAGCAACAGCAGCAGCAACAACAGCGACCGCCACATCTGGTGCACCACGCTGCCAACCACCATCCCGGGCCCGGGGCATGGCGGAGTGCGGCGGCTGCAGCTCACCTCCCTCCCTCCATGGGAGCTTCCAACGGCGGTTTGCTCTATTCGCAGCCGAGCTTCACGGTGAACGGCATGCTGGGCGCAGGAGGGCAGCCGGCTGGGCTGCACCACCACGGCCTGAGGGACGCCCACGATGAGCCACACCATGCAGACCACCACCCGCATCCGCACTCTCACCCACACCAGCAACCGCCCCCGCCACCTCCCCCACAAGGCCCACCGGGCCACCCAGGCGCGCACCACGACCCGCACTCGGACGAGGACACGCCGACCTCAGACGACCTGGAGCAGTTCGCCAAGCAATTCAAGCAGAGGCGGATCAAACTCGGATTTACTCAAGCAGACGTGGGGCTGGCGCTTGGCACCCTGTACGGCAACGTGTTCTCGCAGACCACCATCTGCAGGTTTGAGGCCCTGCAGCTGAGCTTCAAGAACATGTGCAAGCTGAAGCCTTTGTTGAACAAGTGGTTGGAAGAGGCAGACTCATCCTCGGGCAGCCCCACCAGCATAGACAAGATCGCAGCGCAAGGGCGCAAACGGAAAAAGCGGACCTCCATCGAGGTGAGCGTCAAGGGGGCTCTGGAGAGCCATTTCCTCAAATGCCCTAAGCCCTCGGCCCAGGAGATCACCTCCCTCGCGGACAGCTTACAGCTGGAGAAGGAGGTGGTGAGAGTTTGGTTTTGTAACAGGAGACAGAAAGAGAAAAGGATGACCCCTCCCGGAGGGACTCTGCCGGGCGCCGAGGATGTGTATGGGGGTAGTAGGGACACGCCACCACACCACGGGGTGCAGACGCCCGTCCAG

## Design and analysis

Statistical analysis and graphing was completed in Graphpad Prism v6.0. All data is presented as Mean ± Standard Error of the Mean (SEM). As most comparisons comprised only two groups, *t-*tests were the predominant hypothesis-test utilized.

Cohen’s *d* was correspondingly adapted for effect size estimation where of interest, and assumptions screened. Significance was set at *p* < 0.05 per Fisher’s tables, tailed according to statistical-directionality guidelines and corrected for multiple-comparisons. To temper variability, a high-content approach was adapted. Specifically, per sampling theory, we generated, observed, and analyzed as many units of analysis as possible.

Thus, all experiments comprised multiple field, organoid, and human biologic replicates. For multi-cohort replication, samples that were continuously generated for baseline analyses/core phenotype analysis were pooled from across multiple batches, cohorts, and experiments (where possible) to increase data-point capture and maximize power between experiments. To ensure appropriateness, we conducted multiple types of statistical screening to ensure phenotype compliance including a thorough analysis of baseline reproducibility and quality control measures/markers (as previously described in our Methods). Consistent with this, and to promote phenotype representativeness and reproducibility whilst controlling for idiosyncratic differences that may emerge from individual batches, organoids were typically sampled from between 2-6 independent preparations per line that were subsequently pooled. This ensured that organoid quantifications were subjected to high-content sampling to temper human sources of technical (e.g. batch effects) and protocol-derived (e.g. idiosyncrasies) statistical nuisance variance. Indeed, this approach yielded suitable statistical evidence of reproducibility as coefficients of variance between batches were shown to be highly consistent. Additionally, all lines utilized in our rescue studies were pseudorandomly selected for cross-sectional sampling and were not selected based on severity of phenotype or any other qualifying factor. Across immunohistochemistry experiments, up to 12 unique fields from up to 10 cerebral organoid replicates per line were sampled.

This included analysis in up to 21 iPSC lines (see fig. 1) for our core cellular phenotyping experiments. Typically, sampling comprised 2-5 unique observations (VZs or cortical fields) per organoid, and 3-5 organoids were typically sampled per iPSC line. This yielded total sample sizes that often comprised hundreds of observations in sum.

Individual group numbers used for sampling have been provided throughout the manuscript for clarity. Across all experiments no iPSC lines were explicitly/specifically excluded from an analysis. Rather, experiments were conducted in accordance with the production and survival of organoid batches, the relative availability of sections for experiments, as well as indicators of observed power. This cross-sectional sampling approach is consistent with effect sizes and power analyses as well as phenotype replication throughout the manuscript. Cellular quantifications were typically conducted in 5000µm^2^ Regions of Interest (ROI). VZs themselves were defined as progenitor-enriched (SOX2+, PAX6+, NESTIN+) structures that exhibited morphological evidence of a ventricle as well as pseudostratification of neuroepithelium. All cellular analyses were normalized to total levels of the primary cell-type of interest within ROIs (e.g., progenitor death was normalized to quantified progenitor numbers within the same ROI field). Computational analyses were completed, and corrected for false discovery rates, as described above.

## Data availability statement

Proteomics and transcriptomics data used to support the findings of this study have, or will be, deposited in repositories but will be embargoed pending further follow-up analysis/study. However, select –omics data may be accessible following manuscript publication per reasonable request. Code used for transcriptomics data analysis will also be made available by request and/or on Github following manuscript publication.

