## Supplementary Figures and Tables for "BRN2 and PTN unveil multiple neurodevelopmental mechanisms in Schizophrenia patient-derived cerebral organoids"

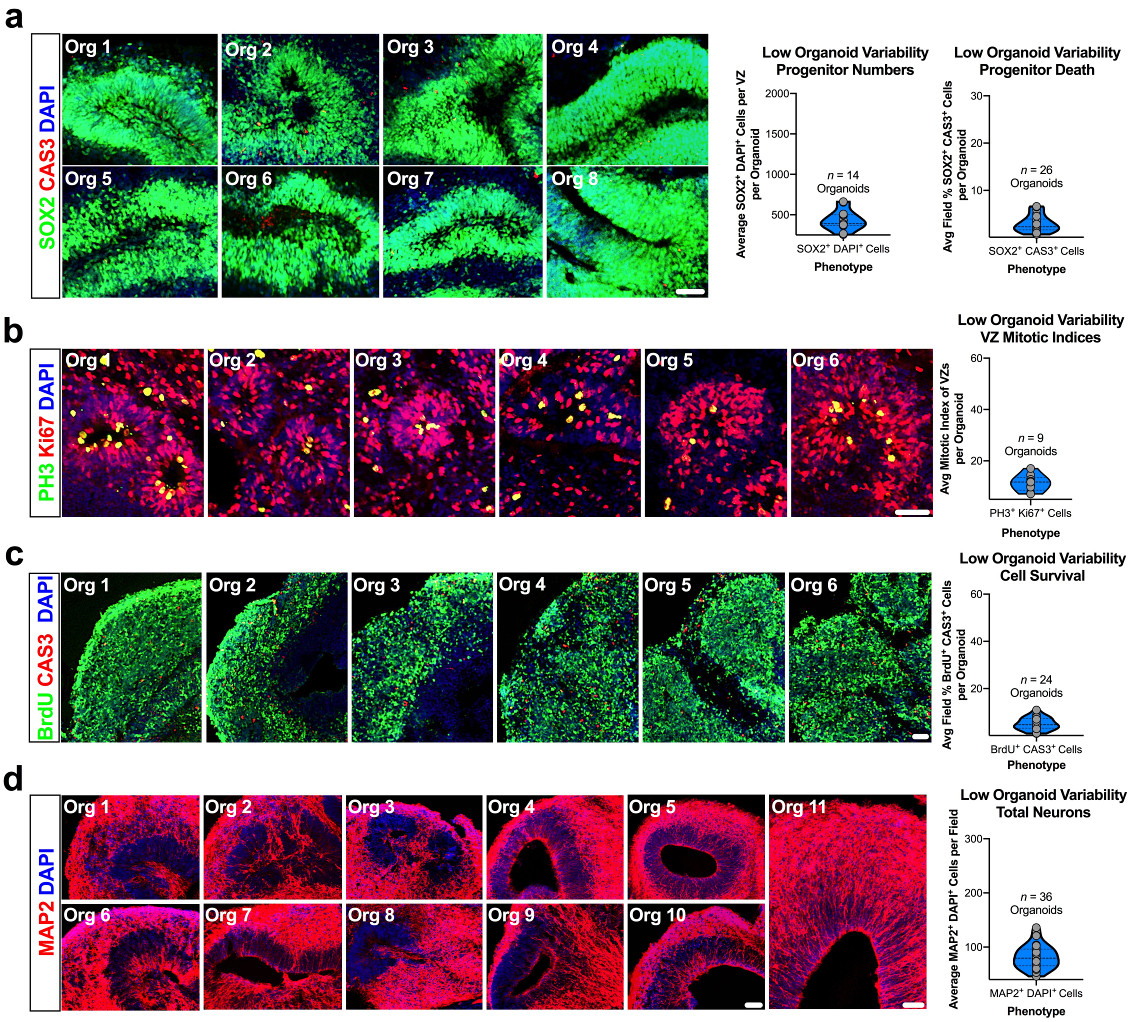


**Figure S1. Unbiased analysis establishes reproducibility of 3D organoids.** Before conducting experiments, we sought to address the variability and reproducibility of our cerebral organoids. We focused on variability of outcome measures that positively reflect organoid health and organization. To do this, we performed multiple non-overlapping quantifications in single organoids, generated an organoid average, and compared this average expression difference to determine variability between organoids. This high-content analysis resulted in hundreds of non-overlapping quantifications from ventricular zones and local fields, derived from a comparatively large number of organoids (typically *n* = 2-5 fields per organoid, *n* = 9-36 organoids, from *n* = 3-5 iPSC lines). All biologics are consistent with those utilized for phenotyping experiments. Overall, this analysis established baseline reproducibility of organoids between- and within-lines, and is consistent with prior unbiased analysis that cerebral organoids hold suitable construct-validity as a model of *in utero* human brain development [1-3].

**a-c, Cerebral organoids exhibit low baseline variability in progenitor pools.** Analysis of multiple regions across *n* = 14 organoids revealed low variability in average progenitor numbers within the ventricular zones of cerebral organoids from healthy subjects. In an even larger analysis of *n* = 26 organoids, variability remained similarly low for progenitor survival between organoids. Cerebral organoids also exhibited reproducible ventricular zone mitotic activity across *n* = 9 individual organoids. To examine new-born cell survival, we adapted the same BrdU pulse-chase assay (24hr pulse, 7d chase) described and utilized in Figures 1 and 5 (see Materials and Methods). Co-staining *n* = 24 individual organoids for BrdU and the cleaved, activated, form of Caspase-3 (CAS3) revealed low and reproducible rates of nascent cell death. Thus, despite differences in morphology, progenitor pool phenotypes were shown to exhibit reproducibility at baseline.

**d, Cerebral organoids exhibit reproducible enrichment of neurons.**
To examine variance in neuron numbers neurons within developing cortical fields, *n* = 36 organoids were stained for MAP2. As can be seen from inset images of cortical fields from 11 different organoids, a robust enrichment of neurons was readily apparent across organoids. When combined, these data established a baseline for cellular variability within cerebral organoids, and confirmed that our organoid cultures exhibited suitably low variability for disease modeling within a morphogen-free 3D macroenvironment.
Ctrl = Control, Scz = Schizophrenia. Each dot on graphs represents the average of a single cerebral organoid. Scale: 60μm.

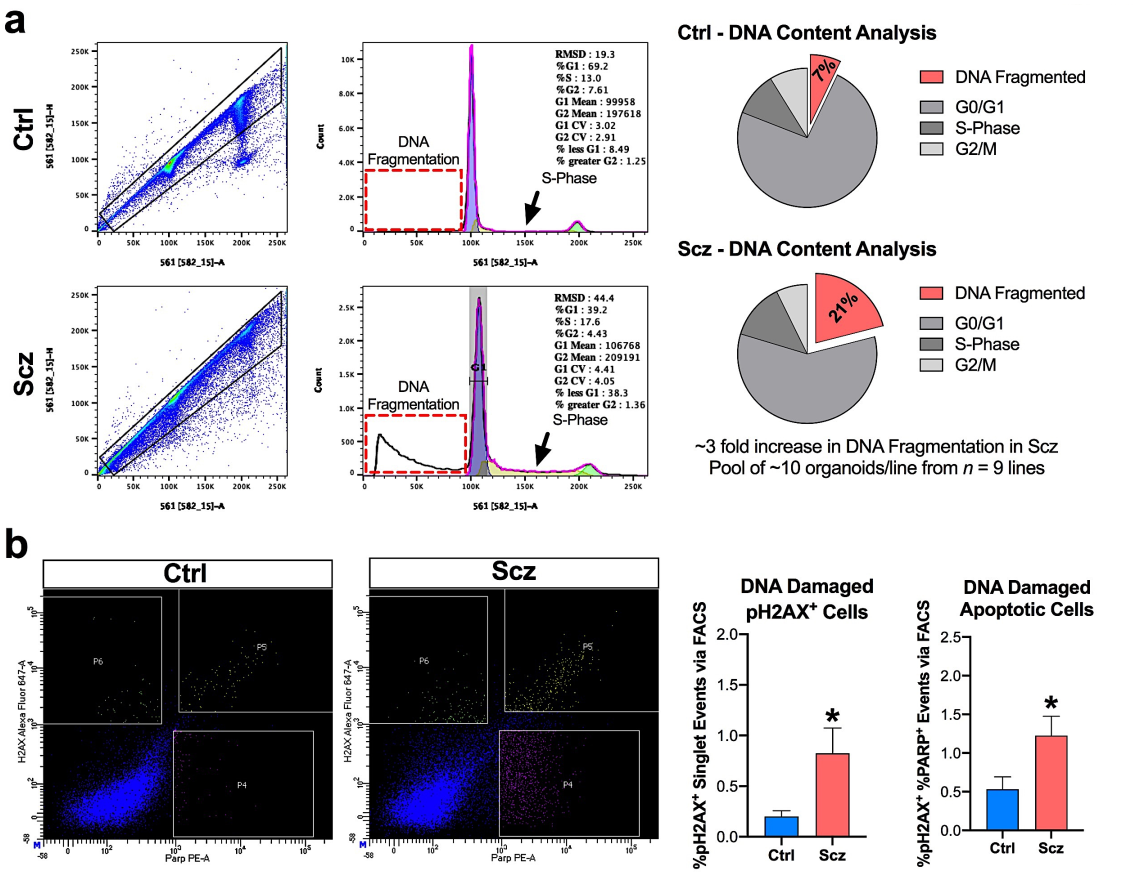


**Figure S2. Validation of cell death in Scz organoids via DNA fragmentation, DNA damage induction, and single-cell PARP+ expression analysis.**
In the main manuscript, we replicated an increase in progenitor cell death via colocalization with cleaved CAS3 (fig. 1, 4, and 5). However, we also sought to validate the increase in apoptosis via an unbiased, DNA-based, methodology to ensure that apoptosis induction in Scz organoids could be orthogonally validated.
**a, Single-cell DNA content supports increased cell death in Scz organoids.** To provide unbiased validation of increased apoptosis in Scz, we subjected a cross-sectional cohort of Ctrl and Scz organoids to a cellular DNA content analysis using Propidium Iodide (PI), flow cytometry, and FlowJo modeling. This allows live and dead cells to be detected based on their DNA content. Notably, when cells undergo apoptosis, DNA becomes fragmented. To reduce sample heterogeneity while promoting power, approximately 10 organoids were pooled per iPSC line (*n* = 9 total lines, 3 Ctrls and 6 Scz), dissociated into a single-cell suspension, and subjected to single-cell DNA content analysis via high-throughput flow cytometry. This analysis revealed that there was an approximately 3-fold increase in DNA fragmentation in Scz organoid samples (~7% in Ctrls vs ~21% in Scz organoids; see red dotted box in panel a). Additionally, Scz organoids also tended to exhibit a slightly elevated proportion of S-phase cells based on cellular DNA content (see black arrows in panel a). When coupled with an increase in DNA fragmentation, an alteration in S-phase may be indicative of a potential DNA-related mechanism that leads to cell cycle abortion and increased death of proliferating progenitors within Scz organoids.

**b, Increased pH2AX recruitment is linked to cell death in Scz organoids.** H2AX is involved in DNA damage [4] and DNA double-strand break repair [5], and is commonly induced during cellular division if DNA damage/breaks have occurred [6]. Phosphorylation of H2AX (pH2AX) indicates the presence of DNA damage machinery [7], and if pH2AX is co-expressed with a cell death factor (e.g. cleaved PARP) then DNA damaged cells are deemed to have been targeted for death. Therefore, based on our neural progenitor cell death and single-cell DNA content analyses, we hypothesized that there would be an increased number of cells exhibiting pH2AX+ as well as pH2AX+ and PARP+ expression within Scz organoids. Organoids were sampled as described above from a pseudorandomly selected cross-section comprising $n = 7$ lines (3 Ctrls and 4 Scz). As expected, we detected an increased percentage of pH2AX+ and pH2AX+/cleaved PARP+ double-positive cells in Scz organoids. This data therefore provides yet further validation of increased cell death in Scz organoids, and suggests that cell death in Scz organoid is related to DNA damage. Graphed data points reflect the average of iPSC lines.
$^*p < 0.05$. Error bars reflect Standard Error of the Mean. Control, Scz: Schizophrenia.

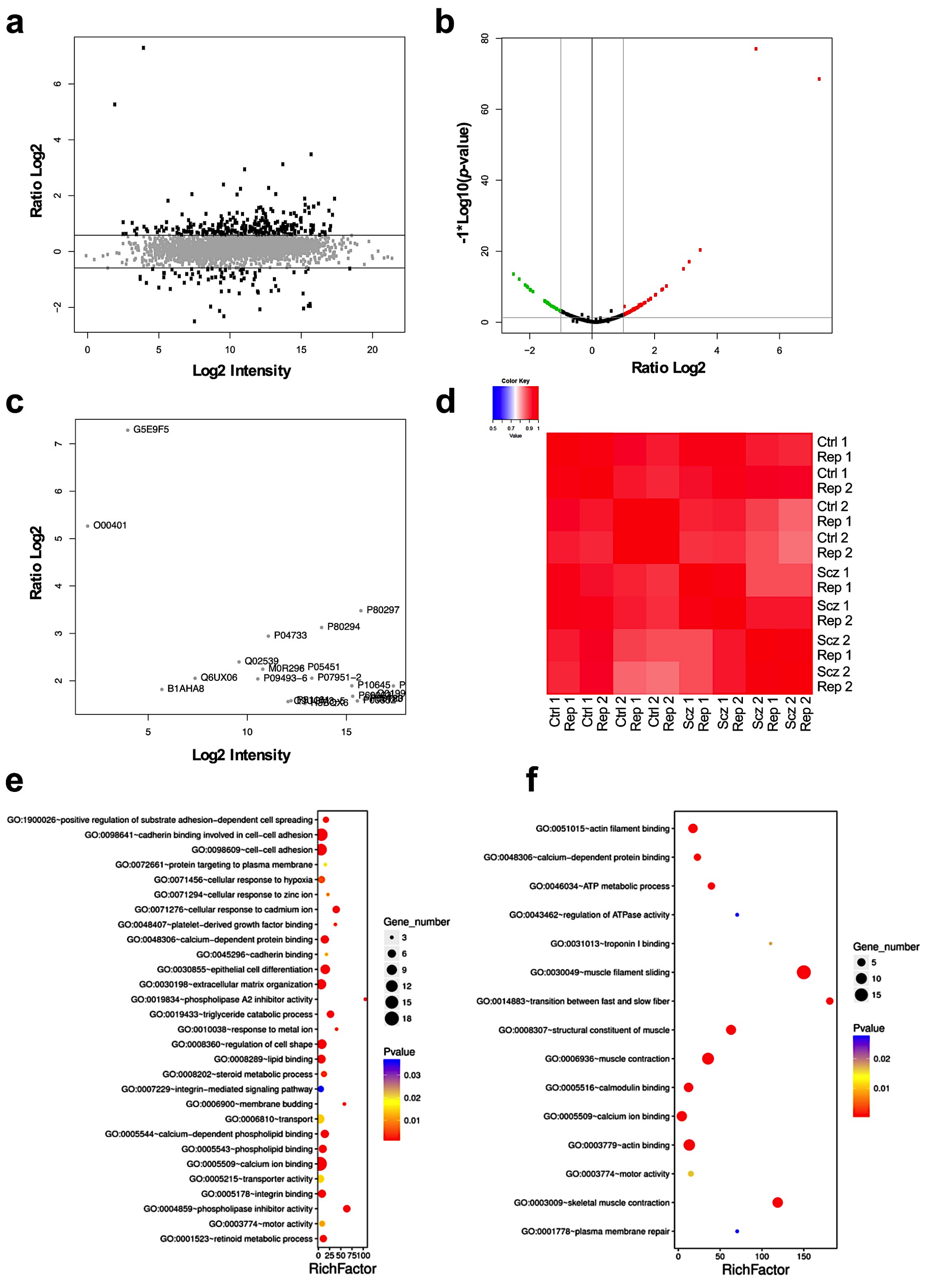


**Figure S3. Further computational analysis of the Scz organoid proteome. a-c, WASL and MPV-17 are novel factors expressed in Scz organoids.** Shown are ratio Log2 against Log2 intensity (a), a volcano plot parsing ratio Log2 by -1*Log10 (*p*-value) (b), and differentially expressed proteins mapped by protein identifiers (c). As can be seen across these panels, differential expression

within the Scz organoid proteome was limited to relatively few targets, suggesting specificity in those proteins that were significantly differentially expressed. The two proteins which with high ratio Log2 values relative to Log2 intensity were WASL (N-WASP) and MPV-17, which were the only two unique proteins detected in Scz organoids. Of note, transcripts for these factors were detected in Ctrl and Scz organoids in single-cell sequencing, but post-translational detection via TMT-LC/MS was restricted to Scz organoid samples. This suggests that N-WASP and MPV17 may be translationally repressed in Ctrl but not Scz organoids. While the functions of MPV17 to Scz remain unclear, the localization of this protein to mitochondria may be indicative of mitochondrial alterations in Scz organoids. Indeed, potential mitochondrial dysfunction in Scz postmortem brains has been reported [8], and mtDNA depletion has been reported in a family with psychosis [9]. Our other novel factor, N-WASP (*WASL*), has a role in cytoskeletal actin assembly that supports dendritic spines [10]. Dendritic spines are typically disrupted in pyramidal layer 3 cells in postmortem Scz tissue [11, 12], where expression of *WASL* also happens to be depleted in postmortem Scz tissue [13]. The post-translational expression of N-WASP (*WASL*) in Scz organoids therefore suggests that this molecule may play a role in early Scz neuropathology in the developing cerebral cortex.
**d, High reproducibility of whole-organoid proteomes across groups.** Shown is a heat map exhibiting all group and replicate samples, which exhibited both 1) global, 2) inter-group, and 3) inter-line reproducibility per coefficient of variance metrics. This once more provides validation of the high reproducibility and low variance of our experimental organoids. Individual protein variation across the proteome was similar across groups and is provided in Figure 2c.
**e-f, Gene ontology and KEGG pathway analysis of the organoid proteome.** Shown are up-regulated (e) and down-regulated (f) protein sets (f; defined as ±1.0 Log2, $p < 0.05$) in Scz patient-derived organoids. Y-axes represent gene ontology (biological process, molecular function, and pathway), and X-axes represents enrichment factor (rich factor = amount of differentially expressed proteins enriched/amount of all proteins in background gene set). Size and color of individual bubbles represents amount of differentially expressed proteins enriched in pathway and enrichment significance, respectively.
Ctrl: Control, Scz: Schizophrenia.

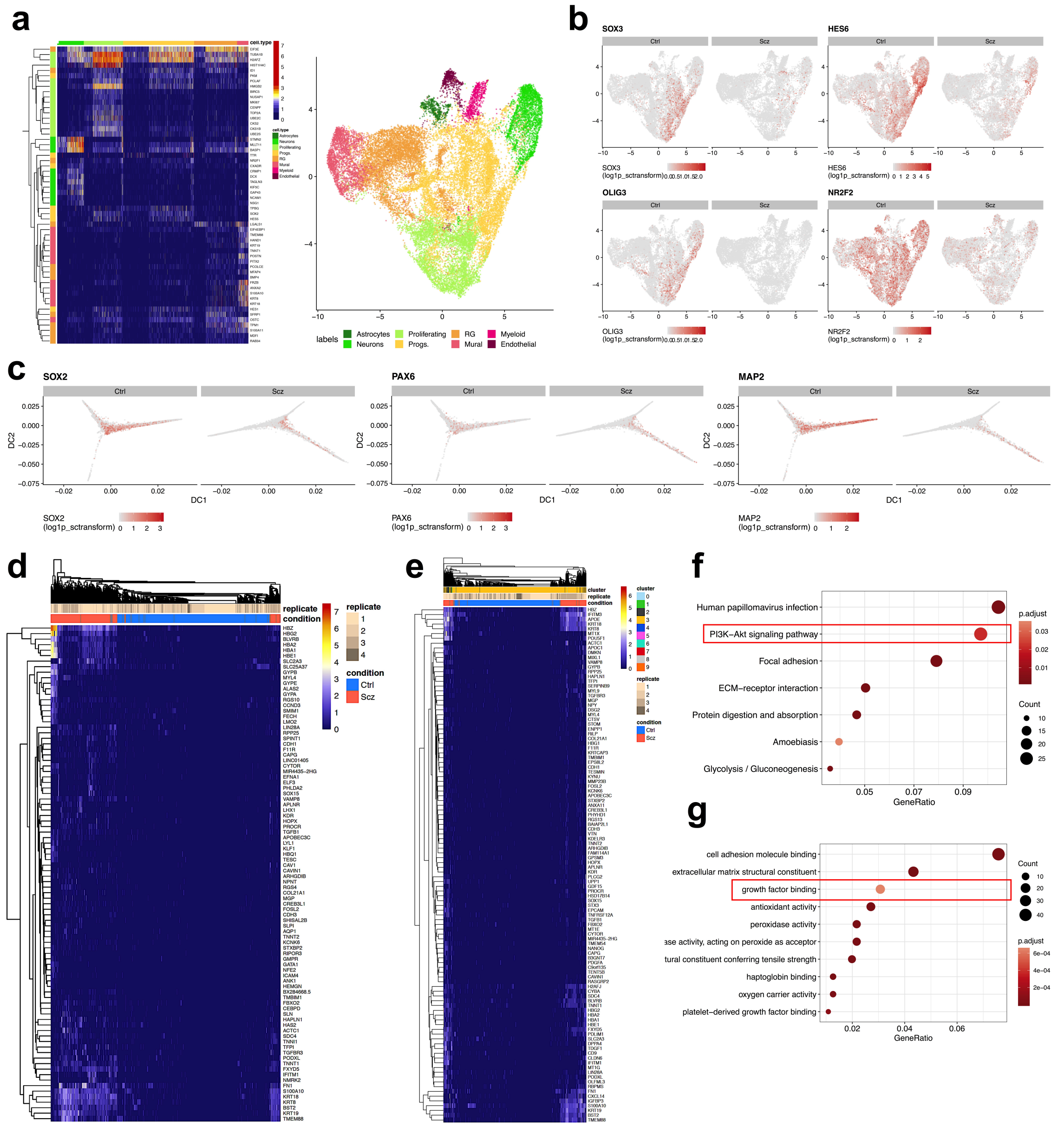


**Figure S4. Further computational analysis of single-cell transcriptomes.**
**a, Heatmap of cell-type clusters across all single-cell transcriptomes.** A heatmap of top differentially expressed factors between cell-type clusters is shown alongside UMAP plots of cell-type clusters for reference.
**b-c, Down-regulated expression of developmental factors in Scz organoids, as well as progenitors and neurons in Scz differentiation trajectories.** Shown are a selection of four noteworthy developmental factors found to be unbiasedly depleted in Scz organoids. Each of these factors have known roles in brain development and/or neuronal maturation. This is consistent with the remodeling of Scz progenitors away from neuron production. Consistent with this, as well as figure 3a-c, shown are additional pseudotime plots for progenitor (*SOX2* and *PAX6*) and neuronal (*MAP2*) markers. These data are consistent with

a depletion of these cell-types in Scz organoids and the remodeling of cell-lineages in Scz organoids, yielding depleted neurogenesis (fig. 1 and fig. 3).

**d-e, Heatmap of upregulated markers in Scz progenitors and neurons.**
In figure 3, we presented cell-specific analysis of top down-regulated factors in Scz single-cell transcriptomes. Here we present heatmaps for top up-regulated factors in Scz progenitors (d) and neurons (e) for evaluation and completeness.

**f-g, Additional plots of pathway analyses of Scz neurons.**
Because individual plots of Scz neuron pathway analysis were not shown in figure 3 due to space considerations, here we present the outcomes of both KEGG (f) and Gene Ontology (g) pathway analyses. Note enrichment for factors which belong to PI3K-Akt signaling (a primary neurotrophin/growth factor signaling cascade) and growth factor binding pathways (red boxes). Enrichment does not depict up- or down-regulation, rather just factor-counts belonging to these biological processes.

Ctrl: Control, Scz: Schizophrenia.

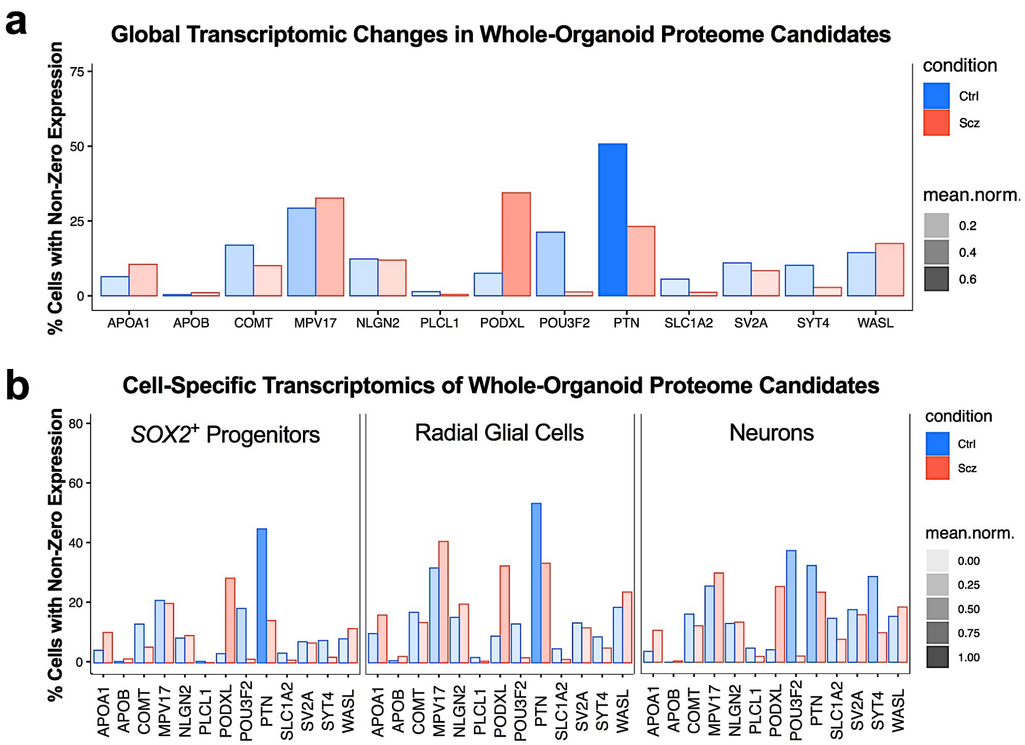


**Figure S5. Cell-specific analysis of proteome targets in Scz organoids.**
**a, Global gene-expression of proteome targets in Ctrl and Scz organoids.** Unbiased analysis of the Scz organoid proteome unveiled that Scz organoids principally differed from Ctrl organoids in the quantity of differentially expressed molecular factors (fig. 2e). This included novel targets that regulate development (e.g. POU-domain transcription factors, e.g. POU3F2/BRN2) or were implicated in genetic risk for Scz in large-scale GWAS and/or meta-analysis (e.g. PTN, PLCL1, COMT, and PODXL, see main text). We cross-referenced expression of these factors in a global analysis of single-cell transcriptomes and found that gene-expression of the forebrain-specific neuronal transcription factor *POU3F2* (BRN2) was down-regulated. Similarly, gene expression of Scz risk candidates *PTN*, *COMT*, and to a lesser extent, *PLCL1*, were also down-regulated. Consistent with proteomics, expression of *PODXL* remained up-regulated in global analysis of Scz transcriptomes.
**b, Cell-specific analysis of proteome targets reveals nuanced expression.** Next, we sought to resolve cell-specific contributions to expression differences as well as gene-expression levels in Scz progenitors and neurons. Thus, we analyzed our scRNA-Seq dataset by cell type (fig. 3h-I and S4). This analysis revealed a striking depletion of *POU3F2* (BRN2) and PTN in Scz progenitor cell-types, as well as a reduction in Scz neurons (as shown in fig. 3). Similarly, *COMT* appeared to be more prominently altered in Scz progenitors, relative to Scz neurons. *PLCL1* expression was more prominently reduced in Scz neurons relative to Scz progenitors. *PODXL* gene-expression was up-regulated in both progenitors and neurons. Ctrl: Control, Scz: Schizophrenia.

**a**

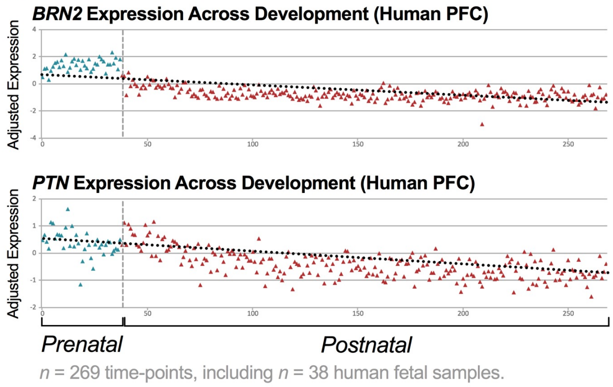


**b**

**SZDB and SZGR Database Hits of Scz Genome-Wide Association**

| Gene | Loci | Index SNP | *p*-value | Loci | Meta-Analysis | Hit | GWAS & Size |
|---|---|---|---|---|---|---|---|
| BRN2 | - | - | - | - | - | - | - |
| PTN | 7:137039644-137028611 | rs3735025 | 7.02e-11 | 55 | ✔ | ✔ | CLOZUK *n* = 35,802 |
| PTN | 7:137039644-137028611 | rs3735025 | 3.28e-09 | 62 | ✘ | ✔ | PGC *n* = 150,064 |

**c**

**SZDB and SZGR Differential Methylation Database Hits**

| Gene | Tissue | ***Probe Location*** | *p*-value | Corrected Value | Study & Size |
|---|---|---|---|---|---|
| BRN2 | Prefrontal Cortex | 6:99280108 | 7.81e-09 | 3.57e-03 | Jaffe et al. *n* = 108 Scz *n = 136* Ctrls |
| BRN2 | Blood | 6:99282348 | 3.34e-02 | - | Kinoshita et al. *n* = 63 Scz *n* = 42 Ctrl |
| PTN | Prefrontal Cortex | 7:136989898 | 1.21e-05 | 1.35e-02 | Wockner et al. *n* = 24 Scz *n* = 24 Ctrls |

**Figure S6. Orthogonal validation of BRN2 and PTN disease relevance via informatic mining of human Scz studies and Scz database repositories.** To provide further insight into the disease relevance of our candidate rescue factors, PTN and BRN2, we consulted large-scale data repositories that have indexed, pooled, or compiled the results of human brain tissue arrays, GWAS, and other commonly employed techniques in Scz association studies.

**a, *BRN2* and *PTN* are enriched during prenatal human cortex development.** To orthogonally validate BRN2 and PTN as embryonic active factors in the developing cortex of humans, we mined the BrainCloud$^{TM}$ (dbGaP Accession: phs000417.v2.p1; [14]) dataset that has quantified gene expression in the human brain across the lifespan. This yielded data congruent with enrichment patterns observed in our organoid cultures, whereby *BRN2* (ProbeID: hHR008743, EntrezID: 5454) and *PTN* (ProbeID: hHC023376, EntrezID: 5764) were expressed in the embryonic human PFC. Additionally, relative to expression patterns across the lifespan, BRN2 and PTN are also enriched during human prenatal development (*n* = 269 human sample time points, including *n* = 38 human fetal samples; dotted lines reflect an approximate line of best fit). These data thus provide orthogonal validation that PTN and BRN2 are transcriptionally enriched during embryonic development of the human cortex.

**b-c, An index SNP in proximity to *PTN* exhibits gene association in Scz.** As discussed in the main manuscript, PTN has been associated with Scz. Here we sought to explore this relationship further by leveraging the Schizophrenia Database (SZDB) [15] and Schizophrenia Gene Resource (SZGR) v2.0 [16, 17]. Both of these references function as multi-omics data repositories that aggregate data from genetic, transcriptome, and epigenetic Scz association studies. Searching these databases and datasets yielded several positive associations of PTN and BRN2 to Scz. This included an index SNP (rs3735025) in proximity to PTN that exhibited genome-wide association within two of the largest Scz cohorts ever studied (total *n* = 35,802 [18] and *n* = 150,064 [19] humans, respectively; see tabulation in b). Additionally, analysis also revealed positive association of both BRN2 and PTN in differential methylation of samples from Scz patients (see [20-22] and tabulation in c for dataset summary). When combined, these data provide orthogonal validation that both of our candidate targets maintain disease relevance to Scz in human association studies.

PFC: Prefrontal cortex.

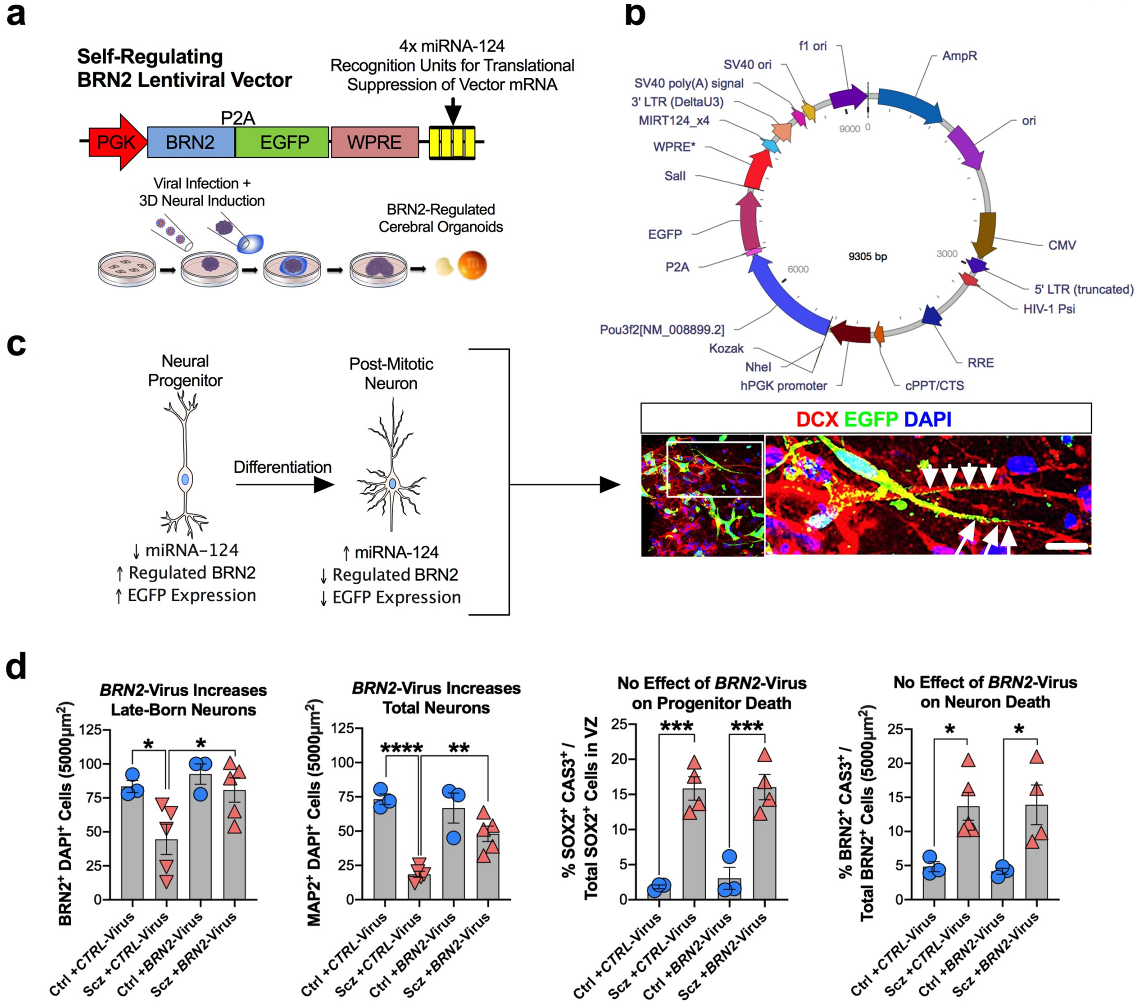


**Figure S7. Supplementary information for BRN2 rescue experiments in Scz patient-derived organoids.**

**a-c, Design of *BRN2*-Virus for rescue experiment in Figure 5.**

Our *BRN2*-Virus is a self-regulating construct designed to be translational suppressed when infected cell-types differentiate into post-mitotic neurons. Thus, sustained overexpression can be induced transiently without continuing to influence neurons. This allowed us to determine in figure 4 that BRN2 influences neuronal differentiation but not survival in Scz organoids. The *BRN2*-Virus construct was adapted from a previously validated construct, but was modified to contain an EGFP reporter construct for visualization (see technical details in Materials and Methods). Our simplified experimental timeline (a) from Figure 5 is provided along with viral construct (b) for comparison and additional details not included in main text for both brevity and simplicity. Also shown is a high-magnification image from within a 3D organoid infected with *BRN2*-Virus that visualizes self-regulated translational suppression via loss of EGFP signal in a differentiating progenitor (arrows) and a new-born neuron (arrowheads; c). As expected, *BRN2*-Virus expression was only observed in progenitors and differentiating cells, but not in mature postmitotic neurons.

**d, Confirmation that *BRN2*-Virus regulates neuron numbers in Scz organoids when data are analyzed by the average of iPSC lines.**

For transparency, here we provide an alternative visualization of *BRN2*-Virus rescue data whereby raw data have been collapsed into the average of iPSC line (each data point here reflects the average of quantifications from an iPSC line). As can be seen in the leftmost panels, *BRN2*-Virus significantly increased neuron numbers when two independent neuronal antigens (BRN2 and MAP2) were utilized for immunohistochemistry and neuron quantifications. Consistent with the data shown in figure 4, neither progenitor nor neuronal survival were rescued by *BRN2*-Virus Scz organoids when the average of iPSC lines was analyzed. Thus, as discussed in the main text, BRN2 was shown to regulate neuron numbers but not cell death phenotypes in Scz organoids.

*$p < 0.05$, **$p < 0.01$, ***$p < 0.001$, ****$p < 0.0001$. Scale: c = 20μm. Error bars reflect Standard Error of the Mean. Ctrl: Control, Scz: Schizophrenia, EGFP = Enhanced green-fluorescent protein.

**a**

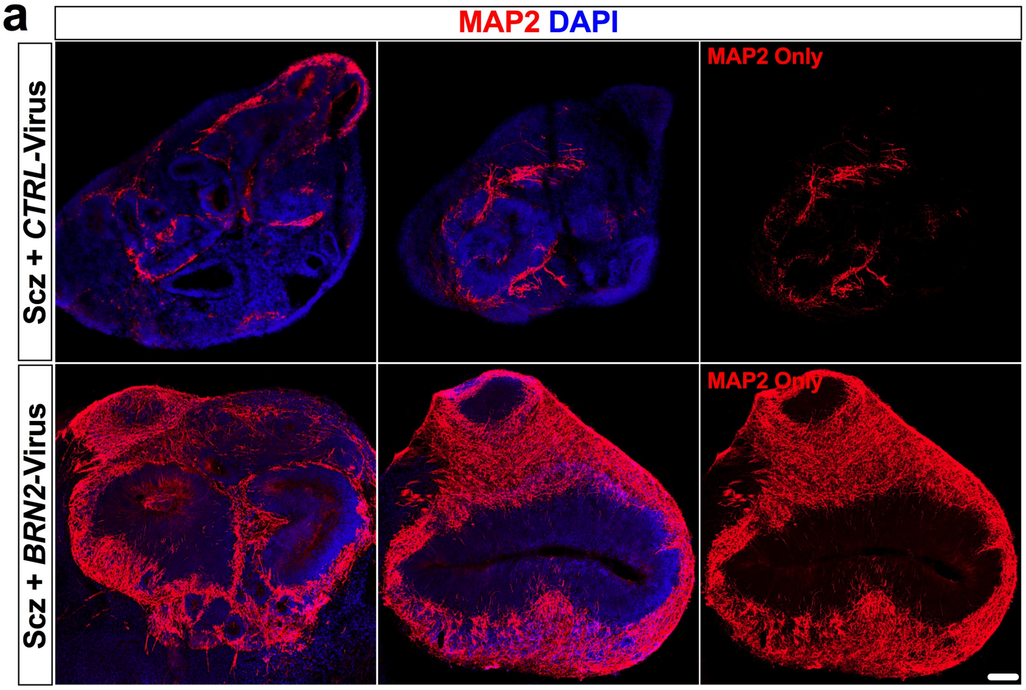


**Figure S8. Enlarged whole-organoid images of *BRN2*-Virus modulation of neuron numbers within Scz organoids.**
To provide additional visualization, we sought to provide additional whole-organoid images of the *BRN2*-Virus rescue effects on neuron numbers in Scz organoids. In the main figures, individual fields were quantified and shown which are representative of both sampling and reported phenotypes. Here we provide whole-organoid images to complement these visualizations, and to exemplify the extent of rescue achieved at the whole-organoid level. Two independent organoids are shown for each treatment group for confirmation of phenotype reproducibility. Note, MAP2 has been presented as an isolated channel for one organoid set for clarity and transparency so that the extent of MAP2 depletion can be observed at baseline in Scz organoids that received *Ctrl*-Virus and the extent of rescue achieved in Scz organoids following infection with *BRN2*-Virus. Note that multiple independent organoids have been selected in the manuscript here to provide numerous independent visualizations of core phenotypes and their rescue in Scz organoids. Scale bar: 60µm.

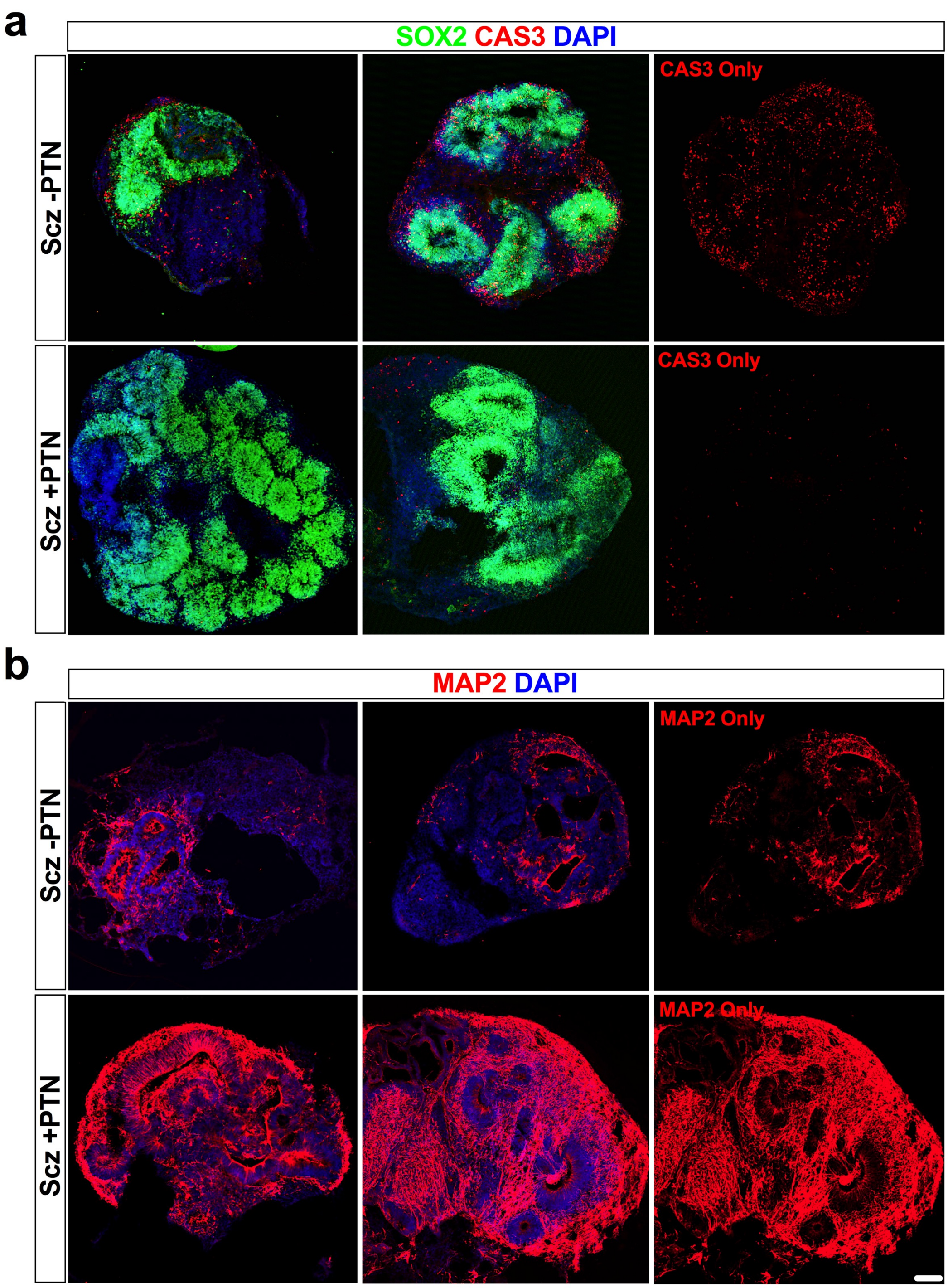


**Figure S9. Whole-organoid images of PTN rescue effects in Scz samples.** To provide additional visualization, we sought to provide additional whole-organoid images of PTN rescue effects in Scz organoids. In the main figures, individual fields were quantified and shown which was representative of both sampling and reported phenotypes. Here we provide whole-organoid images to complement these visualizations, and to exemplify the extent of rescue achieved at the whole-organoid level.

**a, Whole-organoid visualization of progenitor cell death in Scz organoids at baseline and following PTN treatment.**
Two independent organoids are shown for each treatment group for confirmation of phenotype reproducibility. Note, cleaved CAS3 has been presented as an isolated channel for one organoid set for additional transparency.
**b, Whole-organoid visualization of neurons in Scz organoids at baseline and following PTN treatment.**
Once more, two independent organoids are shown for each treatment group for confirmation of phenotype reproducibility. MAP2 has been presented as a split channel for one organoid set for transparency. Note that multiple independent organoids, and fields within them, were shown across the manuscript to provide numerous independent visualizations of core phenotypes and their rescue in Scz organoids. Scale bar: 60 μm

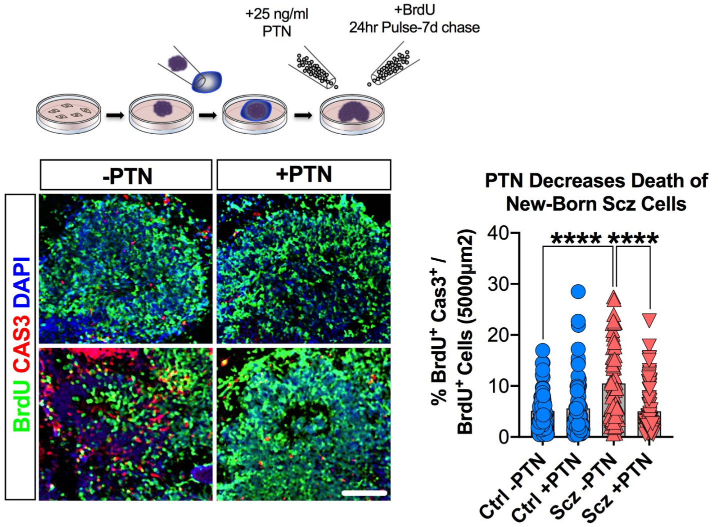


**Figure S10. PTN improves new-born cell survival in Scz organoids.**
In figure 5, we show that PTN-supplemented Scz organoids exhibited increased progenitor survival and neuronal differentiation. In addition to these phenotypes, we also assessed the overall survival of new-born cells labeled with BrdU as per our pulse-chase paradigm (see fig. 1 and 5, as well as our schematic above). PTN-supplemented Scz organoids exhibited significantly decreased new-born cellular death (Vehicle Ctrl $n$ = 71 fields, $n$ = 24 organoids, from $n$ = 4 independent Ctrl iPSC lines, PTN-Supplemented Ctrl $n$ = 95 fields, $n$ = 30 organoids, from $n$ = 4 independent Ctrl iPSC lines; Vehicle Scz $n$ = 69 fields, $n$ = 24 organoids, $n$ = 4 independent Scz iPSC lines, PTN-Supplemented Scz $n$ = 89 fields, $n$ = 26 organoids, from $n$ = 4 independent Scz iPSC lines). All graphed data represent unique, non-overlapping, fields. Together with data in fig. 5, these data indicate that in Scz organoids PTN exerts putative neurotrophic-like effects by promoting cellular survival. Thus, in Scz organoids, reconstitution of PTN as an exogenous morphogen facilitated progenitor survival and neurogenesis that resulted in the rescue of neuron numbers in Scz cortical fields.
****$p < 0.0001$. Scale: 60µm, Error bars reflect Standard Error of the Mean. Ctrl: Control, Scz: Schizophrenia.

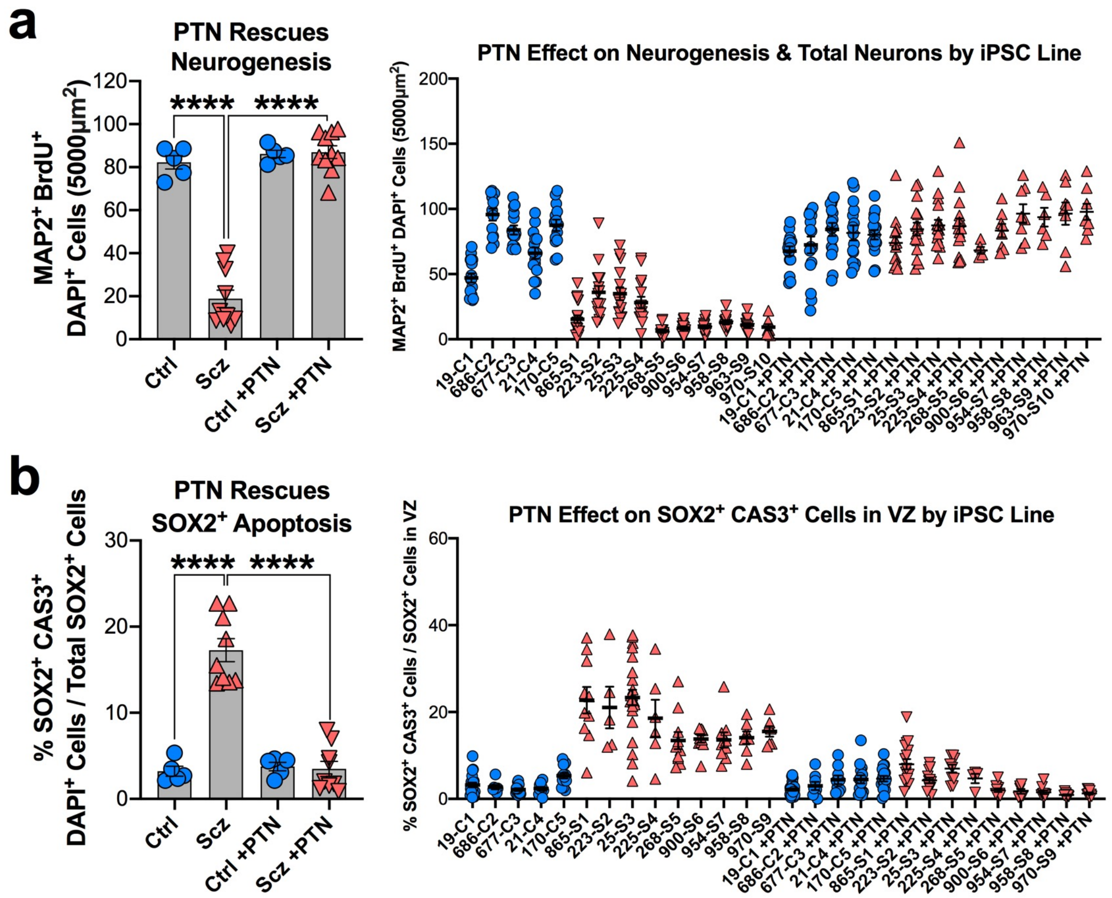


**Figure S11. Analysis of PTN rescue phenotypes by the average of iPSC lines within groups, including graphing of raw data split by iPSC line.** For transparency, visualization of Scz core phenotypes (both at baseline and following PTN treatment) have been provided here as an average of lines (left panels) and raw data for each individual iPSC donor (right panels). Data is provided for all iPSC lines with complete datasets (i.e., both –PTN and +PTN quantifications). This allows the relative consistency of Scz phenotypes to be assessed by each patient iPSC line both with and without PTN treatment. . For patient-split visualizations, note that C1-5 represent biologically unique Ctrl iPSC donors while S1-10 represent individual Scz patients. As can be seen, phenotypes were broadly reproducible across individual lines, and all patient lines included in the PTN rescue experiment exhibited consistency in phenotype via restoration and/or improvement following PTN treatment.

**a, Donor-split analysis of neuronal depletion and death in Scz organoids.** Analysis of neuronal depletion via quantifications of MAP2+ BrdU+ cells revealed that PTN treatment positively regulated neuron numbers. In the iPSC donor-split analysis, each data point represents a single non-overlapping cortical field. Final numbers are provided in the figure legend of the manuscript (fig. 5).

**d, Donor-split analysis of progenitor death in Scz organoids.** Analysis of progenitor death confirmed that Scz organoids exhibited increased SOX2+ cell death in ventricular zones, and that PTN treatment could negatively regulate SOX2+ cell death in Scz organoids. In the iPSC donor-split analysis, each data point represents a unique ventricular zone. Final numbers are provided in the figure legend of the manuscript (fig. 5).

**Supplementary Table 1. Clinical notes of NIMH-deposited Scz iPSC lines.**

| NIH iPSC Scz Line | Sex | Ethnicity | Age of Sampling | Age of Onset | Clinical Notes |
|---|---|---|---|---|---|
| MH0159025 | Male | White | 48 | 41 | Paranoid Scz. Cannabis & alcohol abuse, drug overdose history, father had depression/drug abuse history, one sibling has bipolar disorder, behavioral problems from age 10. |
| MH0159026 | Male | White | 60 | 38 | Persistent auditory hallucinations, lifetime cannabis & alcohol abuse, nicotine addiction, brother had schizophrenia. |
| MH0185223 | Male | White | 26 | - | Episodes of agitation, delusions of persecution, and fear of assassination; at age four mild features of pervasive developmental disorder, Scz/SA/ASD father and sister, brother autistic at age four |
| MH0185225 | Male | White | 23 | - | Paralogical thinking, affective shielding, splitting of affect from content, suspiciousness, SZ/SPD father, anorexic/schizoid sister. |
| MH0200865 | Male | White | 25 | 12 | Childhood onset. Persistent delusional thoughts, persecutory auditory hallucinations, impulsive and hyperactive behavior. Autistic brother. Psychosis scores at baseline (off meds): SAPS = 29, SANS = 49, BPRS24 = 87. |
| MH0217268 | Female | White | 49 | - | - |
| MH0185900 | Male | White | 31 | - | PANSS Total = 71 |

| | | | | | |
|---|---|---|---|---|---|
| MH0185954 | Female | Mixed (Non-Hispanic) | 34 | - | PANSS Total = 100 |
| MH0185958 | Female | White | 56 | - | PANSS Total = 88 |
| MH0185963 | Female | Black | 47 | - | PANSS Total = 71 |
| MH0185970 | Male | White | 52 | - | PANSS Total = 105 |
| MH0185912 | Male | White | 34 | - | PANSS Total = 106 |
| MH0185945 | Male | Hispanic | 58 | - | PANSS Total = 78 |
| MH0185964 | Female | Asian | 43 | - | PANSS Total = 72 |
| MH0185966 | Male | White | 32 | - | PANSS Total = 68 |
| MH0185925 | Male | Mixed (Null) | 64 | - | PANSS Total = 62 |
| MH0185928 | Female | Mixed (Non-Hispanic) | 47 | - | PANSS Total = 88 |

*All controls had no history or reported family history of Scz. Clinical psychometrics and age of onset provided only if known. No further information regarding mixed races was provided from the NIMH beyond the disclaimer that these individuals comprise more than one race. Mixed races were further categorized as “Hispanic” vs. “Non-Hispanic” or “Null” (missing or unknown).

**Supplementary Table 2. Top 10 most abundant proteins in organoids.**

| Protein Description | UniProt Accession | Gene Name | Control Intensity | Scz Intensity |
|---|---|---|---|---|
| Histone H4 | H4_Human | HIST1H4A | 3291771.845 | 2785097.087 |
| Histone H2B | H2B1L_Human | HIST1H2BL | 2592182.54 | 2265873.016 |
| Histone H3.1 | H31_Human | HIST1H3A | 2222150.735 | 1894779.412 |
| Actin, Cytoplasmic 2 | ACTG_Human | ACTG1/ACTG | 1404373.333 | 1417606.667 |
| Histone H2A | A0A0U1RR32_Human | hCG_2039566 | 1286997.041 | 1089911.243 |
| Histone H2A Type 2-B | H2A2B_Human | HIST2H2AB | 1186519.231 | 1061596.154 |
| Tubulin Alpha-1B chain | TBA1B_Human | TUBA1B | 998203.991 | 724672.949 |
| Thymosin beta-4 | TYB4_Human | TMSB4X | 703909.091 | 904840.909 |
| Tubulin Beta-2A Chain | TBB2A_Human | TUBB2A/TUBB2 | 544202.247 | 356531.461 |
| Peptidyl-prolyl Cis-Trans Isomerase A | PPIA_Human | PPIA/CYPA | 426830.303 | 507715.152 |
| Histone H1.5 | H15_Human | HIST1H1B H1F5 | 391176.991 | 330516.593 |
| Vimentin | VIME_Human | VIM | 374522.532 | 458395.923 |

**Supplementary Table 3. Neuronal and disease enrichment in organoids.**

| Pathway Database | Pathway Term | Protein Count | % of Proteome | *p* Value |
|---|---|---|---|---|
| Gene Ontology | GO:0043209: Myelin Sheath | 120 | 3.3632287 | 3.34E-56 |
| Gene Ontology | GO:0043025: Neuronal Cell Body | 94 | 2.634529148 | 6.01E-06 |
| Gene Ontology | GO:0030424: Axon | 72 | 2.01793722 | 3.67E-06 |
| Gene Ontology | Neurogenesis | 69 | 1.933856502 | 6.07E-05 |
| Gene Ontology | GO:0043005: Neuron Projection | 68 | 1.905829596 | 4.42E-04 |
| Gene Ontology | GO:0014069: Postsynaptic Density | 50 | 1.401345291 | 0.008465825 |
| Gene Ontology | Neuropathy | 48 | 1.34529148 | 4.64E-14 |
| Gene Ontology | GO:0007411: Axon Guidance | 43 | 1.205156951 | 0.037049935 |
| Gene Ontology | Epilepsy | 36 | 1.00896861 | 0.002676137 |
| Gene Ontology | GO:0031175: Neuron Projection Development | 34 | 0.952914798 | 0.00195522 |
| Gene Ontology | GO:0007409:Axonogenesis | 28 | 0.784753363 | 0.052012199 |
| Gene Ontology | GO:0010976: Positive Regulation of Neuron Projection Development | 26 | 0.728699552 | 0.048697926 |
| Gene Ontology | GO:0045773: Positive Regulation of Axon Extension | 14 | 0.392376682 | 0.003550075 |

| | | | | |
|---|---|---|---|---|
| Gene Ontology | GO:1990090: Cellular Response to Nerve Growth Factor Stimulus | 14 | 0.392376682 | 0.004992563 |
| KEGG | HSA05016: Huntington's Disease | 95 | 2.662556054 | 4.96E-11 |
| KEGG | HSA05010: Alzheimer's Disease | 84 | 2.35426009 | 3.89E-10 |
| KEGG | HSA05012: Parkinson's Disease | 76 | 2.130044843 | 4.97E-11 |
| KEGG | HSA04530: Tight Junction | 61 | 1.709641256 | 1.59E-05 |
| KEGG | HSA04728: Dopaminergic Synapse | 46 | 1.289237668 | 0.02644444 |
| KEGG | HSA04520: Adherens Junction | 35 | 0.980941704 | 1.36E-04 |
| KEGG | HSA04721: Synaptic Vesicle Cycle | 33 | 0.924887892 | 5.05E-05 |
| KEGG | HSA04720: Long-Term Potentiation | 25 | 0.700672646 | 0.06186726 |

**Supplementary Table 4. Top 10 up-regulated proteins in Scz organoids.**

| Protein Description | UniProt Accession | Gene Name | FC Ratio (RSC) | *p* Value |
|---|---|---|---|---|
| MPV17 | G5E9F5_Human | MPV17 | 156.26 | 2.84E-69 |
| Neural Wiskott-Aldrich Syndrome Protein | WASL_Human | WASL | 38.43 | 9.30E-78 |
| Metallothionein-1X | MT1X_Human | MT1X | 11.149 | 4.27E-21 |
| Metallothionein-1H | MT1H_Human | MT1H | 8.72 | 8.58E-18 |
| Metallothionein-1F | MT1F_Human | MT1F | 7.684 | 9.35E-16 |
| Histone H1a (H1.1) | H11_Human | HIST1H1A | 5.27 | 6.40E-11 |
| Lithostathine-1-Alpha | REG1A_Human | REG1A | 4.842 | 4.69E-10 |
| Uncharacterized Protein | MOR296_Human | - | 4.739 | 7.58E-10 |
| Tropomyosin Beta Chain | TPM2_Human | TPM2 | 4.156 | 1.68E-08 |
| Olfactomedin-4 | OLM4_Human | OLM4 | 4.149 | 1.65E-08 |

**Supplementary Table 5. Top 10 down-regulated proteins in Scz organoids.**

| Protein Description | UniProt Accession | Gene Name | FC Ratio (RSC) | *p* Value |
|---|---|---|---|---|
| Myosin-4 | MYH4_Human | MYH4 | -5.654 | 2.65E-14 |
| Phosphoinositide Phospholipase C-Like 1 | H3BUD4_Human | PLCL1 | -4.977 | 7.04E-13 |
| Myosin-13 | MYH13_Human | MYH13 | -4.363 | 3.19E-11 |
| Troponin C | TNNC1_Human | TNNC1 | -4.195 | 7.47E-11 |
| Myosin-3 | MYH3_Human | MYH3 | -4.111 | 1.41E-10 |
| Troponin C 2 | TNNC2_Human | TNNC2 | -3.917 | 5.99E-10 |
| Histone H2B | U3KQK0_Human | HIST1H2BN | -3.866 | 9.02E-10 |
| Insulin | A6XGL2_Human | INS | -3.845 | 9.63E-10 |
| Myosin Regulatory Light Chain 2 | MLRS_Human | MYLPF | -3.665 | 2.39E-09 |
| Myosin Light Chain 1/3 | MYL1_Human | MYL1 | -2.834 | 8.93E-07 |

**Supplementary Table 6. Top up-regulated GO pathways (proteomics).**

| Pathway Term | Fold Enrichment | *p* Value | Bonferroni Value | False Discovery Rate |
|---|---|---|---|---|
| Extracellular Exosome | 3.65317658 | 1.08E-25 | 2.50E-23 | 1.39E-22 |
| Acetylation | 2.79242172 | 1.17E-14 | 2.60E-12 | 1.49E-11 |
| Extracellular Space | 3.328643311 | 5.98E-09 | 1.39E-06 | 7.68E-06 |
| Annexin* | 69.45219348 | 1.52E-08 | 3.39E-06 | 1.94E-05 |
| Calcium/Phospholipid Binding | 64.82204724 | 2.27E-08 | 5.06E-06 | 2.90E-05 |
| Disease Mutation | 2.542041068 | 3.01E-08 | 6.70E-06 | 3.83E-05 |
| Cadherin Binding Involved in Cell-Cell Adhesion | 6.73508122 | 1.47E-07 | 3.77E-05 | 1.92E-04 |
| Brush Border | 19.28465608 | 1.72E-07 | 4.00E-05 | 2.21E-04 |
| Phosphoprotein | 1.572205851 | 2.27E-07 | 5.06E-05 | 2.90E-04 |
| Collagen Fibril Organization | 24.30603805 | 3.66E-07 | 3.47E-04 | 5.74E-04 |

*Due to repeat protein classifications, Annexin-related pathway categories (namely, IPR001464: Annexin, IPR018252: Annexin Repeat Conserved Site, IPR018502: Annexin Repeat, Repeat: Annexin 1, Repeat: Annexin 3, Repeat: Annexin 4, Repeat: Annexin 2, & SM00335: ANX) were the 5$^{th}$-12$^{th}$ and 14$^{th}$ most significant up-regulated enrichment terms. These categories have been omitted here for repetition, brevity and diversity. Categories ranked by significance.

**Supplementary Table 7. Top down-regulated GO pathways (proteomics).**

| Pathway Term | Fold Enrichment | *p* Value | Bonferroni Value | False Discovery Rate |
|---|---|---|---|---|
| Muscle Protein* | 126.7451613 | 3.65E-38 | 4.70E-36 | 4.24E-35 |
| Muscle Filament Sliding* | 150.0774578 | 3.23E-34 | 1.12E-31 | 4.41E-31 |
| Muscle Myosin Complex* | 189.8333333 | 3.71E-17 | 3.97E-15 | 4.17E-14 |
| Troponin Complex* | 253.1111111 | 9.54E-12 | 1.02E-09 | 1.07E-08 |
| Cardiac Muscle Contraction* | 56.32536688 | 7.44E-11 | 2.58E-08 | 1.02E-07 |
| Actin Binding* | 15.02262774 | 2.12E-09 | 2.74E-07 | 2.47E-06 |
| Domain: IQ | 36.54462659 | 5.64E-07 | 4.45E-04 | 8.64E-04 |
| Disease Mutation | 3.228392157 | 7.20E-07 | 9.29E-05 | 8.37E-04 |
| Methylation | 5.233566434 | 1.36E-06 | 1.76E-04 | 0.001582906 |
| Calmodulin Binding | 17.23289474 | 2.83E-06 | 3.65E-04 | 0.003295404 |

*Due to repeat protein classifications, largely derived from pathways with overlap involving myosin-, troponin- and actin-related proteins, several GO categories with higher significance than Domain:IQ, Disease Mutation, Methylation and Calmodulin Binding were omitted. These included protein classifications consisting of GO:0003009: Skeletal Muscle Contraction, Thick Filament, GO:0032982: Myosin Filament, GO:0006936: Muscle Contraction, Myosin, GO:0008307: Structural Constituent of Muscle, IPR004009: Myosin N-Terminal SH3-Like, GO:0030017: Sarcomere, IPR002928: Myosin Tail, IPR027401: Myosin-Like IQ Motif, Motor Proteins, GO:0000146: Microfilament Motor Activity, GO:0003779: Actin Binding, Region of Interest: Actin Binding, Domain: Myosin Head-Like, GO:0006937: Regulation of Muscle Contraction, IPR001609: Myosin Head, Motor Domain, SM00242: MYSc, IPR001978: Troponin, GO:0114883: Transition Between Fast and Slow, GO:0030017: Z Disc, GO:0030016: Myofibril, & GO:0051015: Actin Filament Binding. These GO categories largely reflect the early classification of myosins and other related proteins to muscle pathways and protein classification sets, despite being found in other organs, tissues and cell groups of non-muscular origin. Categories ranked by significance.
